# Resolving fate and transcriptome of hematopoietic stem cell clones

**DOI:** 10.1101/2020.03.25.008433

**Authors:** Weike Pei, Fuwei Shang, Xi Wang, Ann-Kathrin Fanti, Alessandro Greco, Katrin Busch, Kay Klapproth, Qin Zhang, Claudia Quedenau, Sascha Sauer, Thorsten B. Feyerabend, Thomas Höfer, Hans-Reimer Rodewald

## Abstract

Adult bone marrow harbors a mosaic of hematopoietic stem cell (HSC) clones of embryonic origin, and recent work suggests that such clones may have coherent lineage fates. To probe under physiological conditions whether HSC clones with different fates are transcriptionally distinct, we developed *PolyloxExpress* – a Cre recombinase-dependent DNA substrate for in situ barcoding that allows parallel readout of barcodes and transcriptomes in single cells. We describe differentiation-inactive, multilineage and lineage-restricted HSC clones, find that they reside in distinct regions of the transcriptional landscape of hematopoiesis, and identify corresponding gene signatures. All clone types contain proliferating HSCs, indicating that differentiation-inactive HSCs can undergo symmetric self-renewal. Our work establishes an approach for studying determinants of stem cell fate in vivo and provides molecular evidence for fate coherence of HSC clones.

---

Hematopoietic stem cell (HSC) emerge in mice from around embryonic day (E) 9.5 onwards. HSC progenitors give rise to definitive HSC which expand, migrate to the fetal liver and colonize the bone marrow around birth, where they persist and function lifelong. In the bone marrow, HSC generate at low rate multipotent progenitors (MPP) (*1-5*); these progenitors self-renew and play a major role for daily hematopoiesis in the adult (*1, 6-8*). Transposon-tagging of HSC in situ has indicated the existence of multilineage HSC clones giving rise to lymphoid and myelo-erythroid output, myelo-erythroid-restricted HSC clones lacking lymphoid output, and inactive HSC clones that do not produce differentiated output (*6, 9*). Single lineage-fate, megakaryocyte-producing HSC have also been reported (*9, 10*). The existence of coherent lineage fates of HSC in a clone is further supported by in situ color-coding of HSC, followed by transplantation (*11*). To understand the development of HSC clones, and the fates emerging from these clones, we previously developed Cre-driven high-resolution barcoding of single HSC in the fetus, which allows tracing embryonically-derived HSC clones to the adult bone marrow and their output (*12*). Barcode induction starting at E9.5, and lasting until about E11.5 (*13*), covering HSC emergence and expansion (*14*), labels around 90% of all adult HSC in the bone marrow with barcodes (*12*). HSC clones persisting in the adult were found to be inactive (defined as no output of erythroid, myeloid or lymphoid cells), myelo-erythroid-restricted, or multilineage (*12*). The collective findings by others and us raise the question of whether the coherent lineage fate of HSC clone is transcriptionally determined. To address this question, we have devised an endogenous RNA barcoding system, *PolyloxExpress*, that allows the joint read-out of barcodes and transcriptomes in single cells. We use *PolyloxExpress* to study together fate (via comparison of barcodes in HSC and mature lineages), and transcriptome (via single-cell RNA sequencing and barcode matching) of individual HSC clones in mice.

## *PolyloxExpress* integrates lineage information and transcriptome in single cells

To read barcodes for fate analysis together with transcriptomes, we developed the *Rosa26*^*PolyloxExpress*^ mouse in which barcodes are generated in genomic DNA and transcribed as mRNA. The original *Polylox* DNA substrate cassette (*12*), composed of nine unique DNA blocks with ten intervening *loxP* sites in alternating orientations, was targeted in ES cells into the *Rosa26* locus downstream from a tdTomato fluorescence reporter (Fig. 1A; fig. S1A, B). The tdTomato reporter revealed mRNA expression from this engineered locus (fig. S1C). Cre-driven random deletion or inversion of *loxP* site-flanked DNA blocks in the *Polylox* DNA substrate results in the generation of DNA barcodes (*12*) (Fig. 1A).

**Fig. 1.**
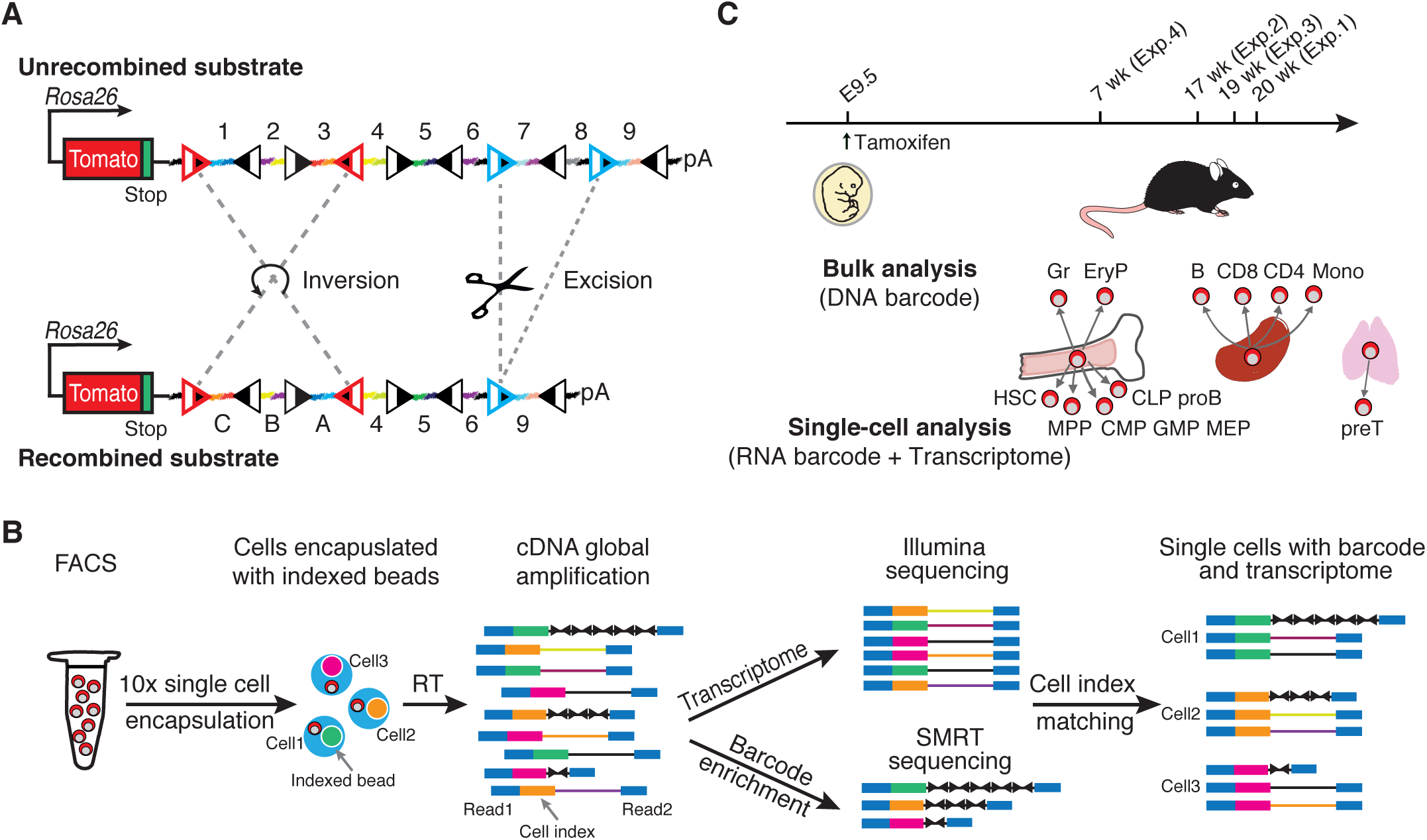
*PolyloxExpress* for simultaneous lineage tracing and transcriptome profiling at single-cell resolution. **(A)** Schematic diagram of the *PolyloxExpress* barcoding system. The *PolyloxExpress* cassette contains a tdTomato reporter and the *Polylox* substrate. In the unrecombined *Polylox* substrate, unique barcode segments (colored linkers, abbreviated 1–9 and A-I when inverted) are interspaced by ten *loxP* sites (triangles). Upon Cre activation, barcode segments are rearranged by excision (blue) or inversion (red) (*12*). Driven by the *Rosa26* promoter, tdTomato and *PolyloxExpress* barcodes followed by a poly-Adenylation signal (pA) site are transcribed as mRNA. **(B)** *Polylox* barcode detection and transcriptome profiling using single-cell RNA sequencing. Cells purified by fluorescence-activated cell sorting (FACS) were captured by the 10x Genomics microfluidic system. After reverse transcription (RT) in droplets, pooled cDNA was amplified and split into two aliquots for parallel transcriptome library preparation (upper arm) and barcode enrichment (lower arm). *Polylox* barcode and transcriptome were integrated by cell-specific index matching. **(C)** Schematic diagram of HSC fate mapping *in vivo*. Barcodes were induced in HSC precursors in *Tie2*^*MeriCreMer/+*^ *Rosa26*^*PolyloxExpress/+*^ embryos. In adult mice at indicated time points, hematopoietic stem and progenitor cells were analyzed by single-cell RNA sequencing for transcriptomes and barcodes, and the indicated peripheral cells by DNA sequencing for barcodes. Abbreviations: Gr, granulocytes; EryP, erythroid-progenitors stage II/III; Mono, Monocytes; HSC, hematopoietic stem cell; MPP, multipotent progenitor; CMP, common myeloid progenitor; GMP, granulocyte-monocyte progenitor; MEP, megakaryocyte-erythrocyte progenitor; CLP, common lymphoid progenitor.

In bulk cultures of MeriCreMer (tamoxifen-inducible Cre) (*15*) transfected *Rosa26*^*PolyloxExpress/+*^ ES cells, DNA and RNA barcodes were induced by hydroxy-tamoxifen with parallel kinetics (fig. S2A, B). DNA and RNA barcodes were identical in single ES cell clones (fig. S2C, D). To identify barcode and transcriptome from single hematopoietic cells, we developed a workflow composed of (1) single-cell encapsulation of flow cytometer-sorted stem and progenitor cells with indexed beads on the 10x Genomics platform, (2) cDNA library preparation and amplification, and (3) splitting of these libraries for whole transcriptome analysis via Illumina sequencing, and targeted *Polylox* amplification and SMRT sequencing of barcodes. Cell indices were retrieved from both sequencing arms (Illumina and SMRT) to match barcode and transcriptome for single cells (Fig. 1B; fig. S3A, B shown for ES cells). *Polylox* reads were filtered for correct, complete and unambiguous barcodes (fig. S3C).

For in vivo experiments, we generated *Rosa26*^*PolyloxExpress/+*^ (termed *Rosa26*^*PolyloxExpress*^) mice. Barcode induction in *Rosa26*^*PolyloxExpress*^ mice was strictly dependent on Cre activity (fig. S4). To induce barcodes in HSC clones at the time of their generation, we treated *Tie2*^*MeriCreMer*^*Rosa26*^*PolyloxExpress*^ mice bearing inducible Cre recombinase in the *Tie2* locus in utero at E9.5 with tamoxifen, which transiently activates Cre in HSC progenitors (*1, 12*) (Fig. 1C). Counting the number of different barcodes in all sampled hematopoietic cells per adult mouse (see Materials and Methods) revealed a barcode diversity of 740 ± 252 (mean ± standard deviation [SD] from Exp. 1-4, table S1) different barcodes, with 88.3% (±7.6; mean ± SD, table S1) of cells being barcoded. This number is in keeping with estimated numbers of several hundred embryonic HSC progenitors (*16*), suggesting that barcode induction in *Tie2*^*MeriCreMer*^*Rosa26*^*PolyloxExpress*^ mice at E9.5 covers emerging HSC quantitatively. When restricting the analysis to HSC and progenitors retrieved collectively from all major adult mouse bones, on average 377 ± 128 (mean ± SD from Exp. 1 - 3, table S1) different barcodes were found. Hence, about 50% of the number of embryonically induced barcodes were retrieved in HSC and progenitors in the bone marrow, suggesting that these barcodes represent HSC clones that colonized the bone marrow and persisted there.

To reveal HSC fates, mice (Exp. 1 - 4) were analyzed between 7 and 20 weeks after birth (Fig. 1C) (table S2). We sorted HSC, MPP, common myeloid progenitors (CMP), granulocyte-monocyte progenitors (GMP), megakaryocyte-erythrocyte progenitors (MEP), common lymphoid progenitors (CLP), pro B cells, pre T cells, CD4 and CD8 T cells and B cells, granulocytes, monocytes and erythrocyte progenitors (EryP) (Fig. 1C; cells were gated and sorted according to published phenotypes (*12*), and fig. S5). Single HSC and progenitors were analyzed for barcodes and whole transcriptome. Barcode correlations between sample repeats for single-cells (fig. S6A) or bulk populations (fig. S6B) revealed equally robust barcode recovery from progenitors and mature cells. Moreover, barcodes identified by single-cell RNA sequencing were also found in the bulk analysis of the same population, with very few exceptions (fig. S6C). On average, 18.8% of HSC and progenitor barcodes, and 20.5% of mature cell barcodes were recovered only in one of two samples, suggesting that the probability to completely miss a low-frequency barcode in any given population (false-negative rate) is a mere 4%. The correlation patterns of barcode frequencies in the peripheral cell populations (fig. S7A) were consistent with a major split between myelo-erythroid and lymphoid development (fig. S7B).

## Projection of HSC clones onto the transcriptional landscape

We defined individual HSC clones by focusing on *Polylox* barcodes with low generation probability (*P*_gen_) as described previously (*12*); we chose a threshold of *P*_gen_ < 5 × 10^−4^, ensuring that more than 90% of rare barcodes were induced only once in HSC progenitors (*17*). Across the four experiments (Fig. 1C), this yielded 91 HSC clones that had the following fate patterns: (1) Inactive HSC clones in which barcodes found in HSC were absent from all peripheral lineages; inactive HSC clones hence lacked lineage output; (2) Myleo-erythroid-restricted, or biased, HSC clones harboring barcodes that were found in myeloid cells and erythrocytes, but only marginally present or absent in lymphoid lineages; (3) Multilineage HSC clones in which barcodes found in HSC were present in all sampled lineages; (4) a small number of clones with low barcode frequencies across several but not all myelo-erythroid and lymphoid lineages were not classified (Fig. 2A; fig. S7C, D).

**Fig. 2.**
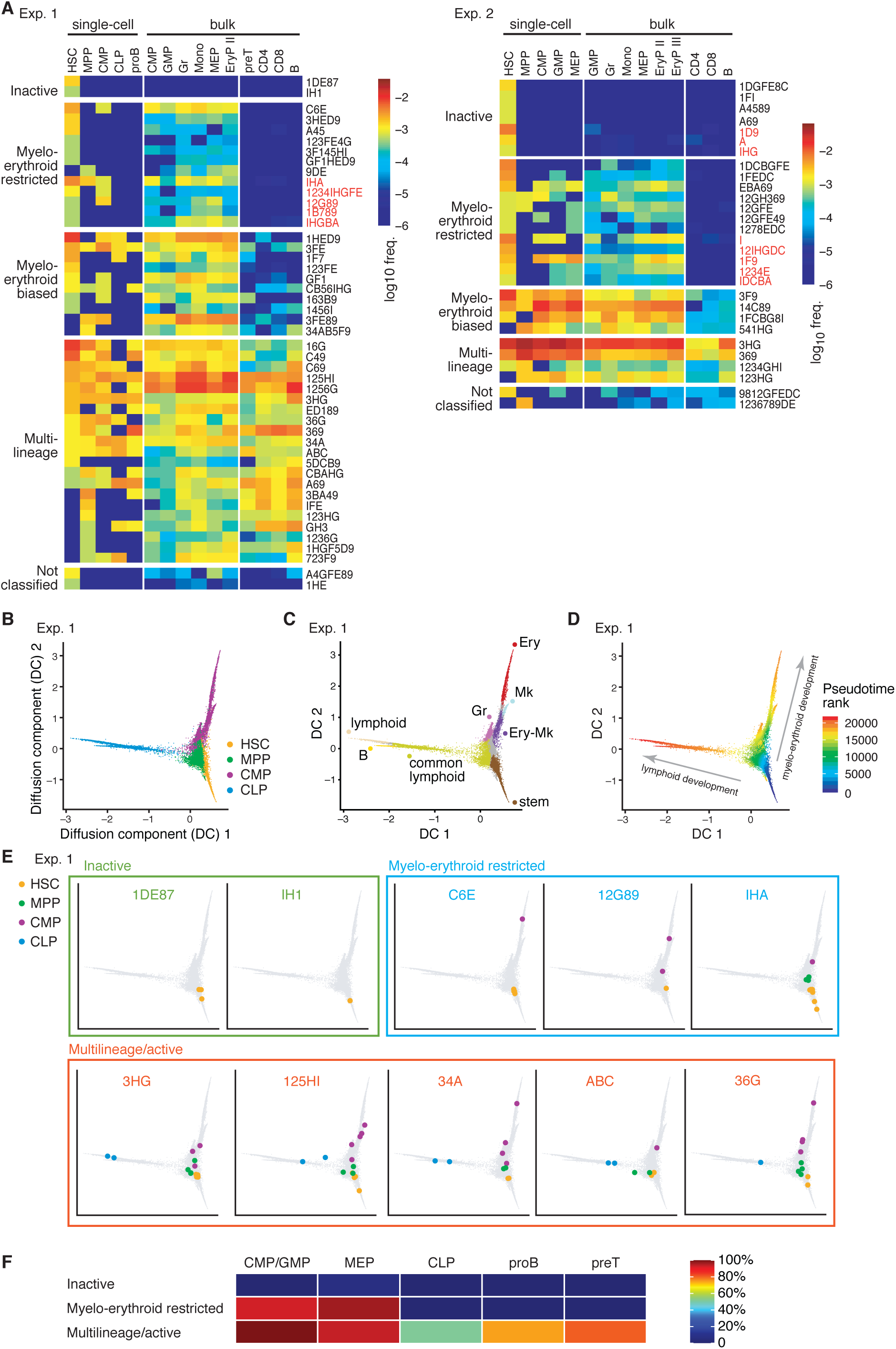
Projection of fate-defined HSC clones onto the hematopoietic transcriptional landscape. (**A**) Heatmaps of barcode read counts in HSC (first lane), MPP (second lane), indicated progenitors, and erythroid, myeloid and lymphoid lineages. Based on the presence of HSC barcodes in peripheral lineages, HSC clones were classified into inactive, myelo-erythroid-restricted, myelo-erythroid biased and multilineage. Color scale indicates barcode frequencies. Rare barcodes (*P*_gen_ < 5 × 10^−4^) are shown in black; barcodes with *P*_gen_ > 5 × 10^−4^ (red) showing no output (inactive) or myelo-erythroid-restricted output are included. (**B**) Transcriptional landscape of phenotypically defined cell populations (HSC, MPP, CMP, and CLP) in Exp. 1 reconstructed using the diffusion map. Each dot represents a single cell of the four cell populations. (**C**) Identification of branches in the transcriptional landscape from Exp. 1 using diffusion pseudotime. Lineage branches are color-coded and labeled according to marker gene expression. (**D**) Pseudo-temporal ordering of cells according to pseudotime rank. (**E**) Projection of fate-defined hematopoietic stem and progenitor cells onto the transcriptional landscape. In individual plots, cells derived from the indicated HSC clone, colored according to their phenotypically defined stem and progenitor stages, are located on the landscape (grey background). Examples of inactive (green frame), myelo-erythroid-restricted (blue frame), and multilineage clones (red frame), all defined in (A), are shown. (**F**) Heatmap showing which fraction of HSC clones of a given fate category producing common myeloid (CMP/GMP), megakaryocyte-erythroid (MEP), common lymphoid (CLP), B cell (proB) or T cell (preT) progenitors.

Next, we analyzed the single-cell transcriptomes of sorted HSC and progenitor cells. (Fig. 2B, fig. S8A, E, I). To reveal the transcriptional landscape of HSC differentiation, we used the diffusion map (*18*). This unsupervised computational technique robustly yields putative differentiation pathways through neighboring cells in transcriptional space (spanned by the expression of the approximately 1000 highly variable genes) and, indeed, identified correctly the trajectories of known stem cell and differentiation markers (fig. S9). Branches were defined using the diffusion-pseudotime algorithm (*19*) and annotated using lineage-specific marker genes (table S3, Materials and Methods), revealing major branching into lymphoid versus myeloid, erythroid and megakaryocyte lineages (Fig. 2C; fig. S8B, F, J). The pseudo-time rank measures position of a cell in this landscape, starting from tip HSC in the common trunk towards the diverse types of progenitors (Fig. 2D; fig. S8C, G, K).

To locate fate-defined HSC clones on the transcriptional landscape, we sorted HSC clones according to their fates (inactive, myelo-erythroid-restricted, multilineage). We found that HSC belonging to a fate category were transcriptionally more similar with each other on the landscape than randomly drawn HSC (fig. S10). This finding suggests that coherent fates of HSC in a clone are rooted in similar transcriptomes.

The type of short-lived progenitor cells generated by an HSC clone indicates its current production of differentiated cells. In the transcriptional landscape, inactive HSC clones had no progenitor cells in lineage branches, which is consistent with the absence of peripheral products (Inactive: Fig. 2E, green panel; fig. S8D, H, L, green panels). Myelo-erythroid HSC clones, including their products, resided in the HSC and MPP territories and encompassed further differentiated progenitors along myelo-erythroid branches (Myelo-erythroid-restricted: Fig. 2E, blue panel; fig. S8D, H, L, blue panels). Cells belonging to multilineage HSC clones were found at HSC and MPP stages and contained both lymphoid and myelo-erythroid progenitors (Multilineage: Fig. 2E, orange panel; fig. S8D, H, L, orange panel). Specifically, we never found a lymphoid progenitor made by a myelo-erythroid-restricted HSC clone or other violations of the HSC clone fate (Fig. 2F). Thus, the distribution of progenitor cells is, without exception, consistent with the fates of HSC clones defined by barcodes in peripheral cells.

## Molecular hallmarks of active and inactive HSC clones

To elucidate molecular differences between inactive and active multilineage HSC, we resolved their positions in the trunk of the transcriptional landscape (Fig. 3A). Inactive HSC in individual experiments were closer to the root of the landscape compared to active HSC clones (Fig. 3B). We asked whether these positions correlated with previously described HSC subpopulations. Our sorted, fate- and transcriptome-analyzed HSC contained both phenotypic long-term (LT) (Lin^-^ Kit^+^Sca1^+^CD150^+^CD48^-^) and short-term (ST) (Lin^-^Kit^+^Sca1^+^CD150^-^CD48^-^) HSC (*20*). These categories were previously defined based on reconstitution duration after cell transplantation, whereas fate mapping showed that ST-HSC are long-term, self-renewing contributors to physiological hematopoiesis with a residence time of 330 days (*1*). Analyzing the expression of LT and ST specific genes (table S4; Materials and Methods), inactive HSC scored more highly for the LT gene expression signature than active HSC, which expressed higher levels of the ST signature (Fig. 3C). In terms of cell frequencies, inactive HSC were enriched for LT-HSC, while active HSC for ST-HSC (Fig. 3D). Nevertheless, there were both active and inactive HSC with LT and ST signatures.

**Fig. 3.**
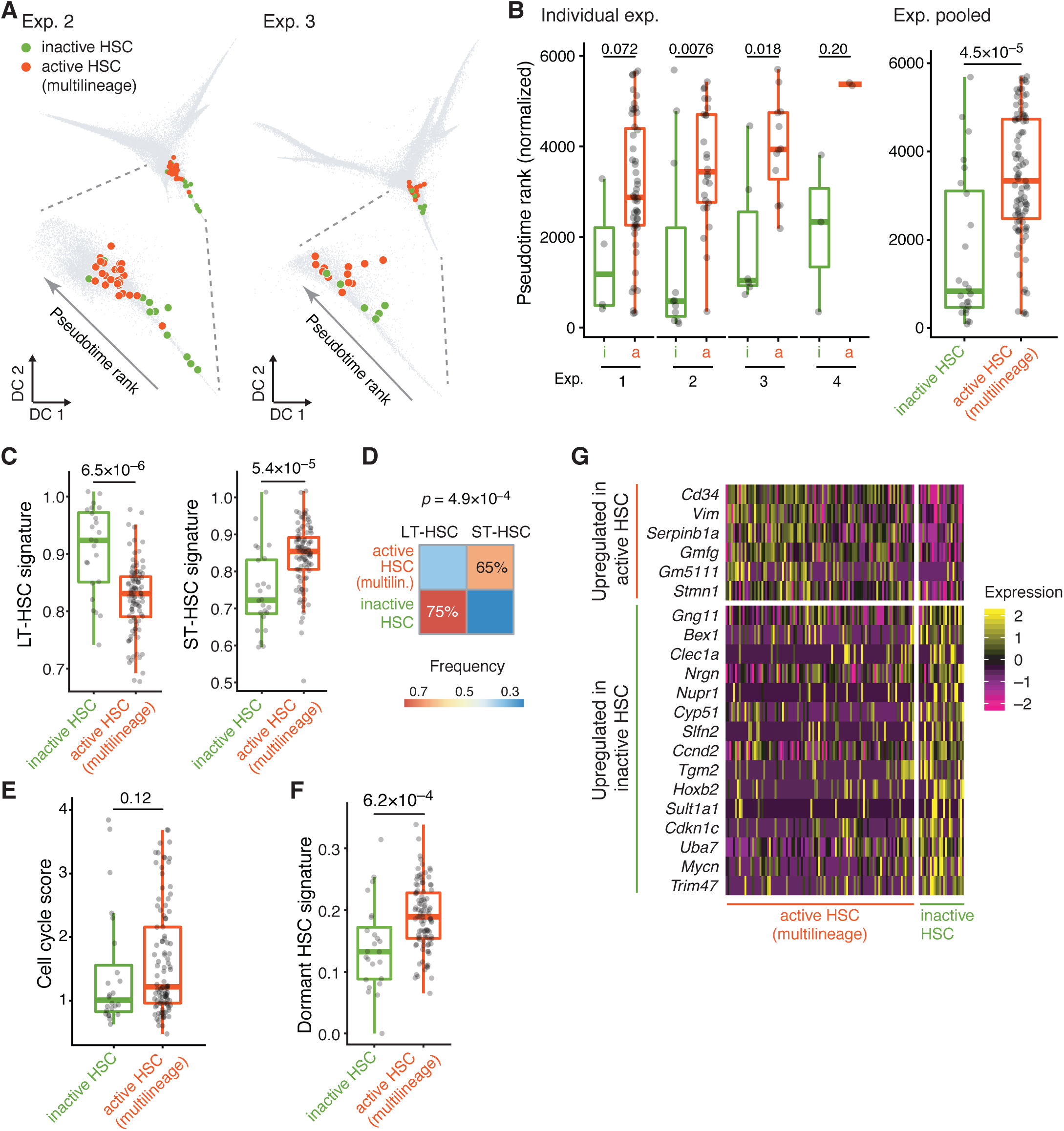
Transcriptomic characterization of inactive and active HSC clones. (**A**) Projection of inactive and active HSC on the transcriptional landscape (as in Fig. 2E) for Exp. 2 and 3. A magnified view of the stem cell trunk is shown for each dataset. Multilineage HSC clones were taken as active clones [n = 11 (Exp. 2) and 6 (Exp. 3) inactive HSC; n = 29 (Exp. 2) and 13 (Exp. 3) active HSC]. **(B)** Statistical comparison of the pseudotime ranks between inactive and active (multilineage) HSC (for individual experiments: n = 4, 11, 6 and 3 inactive HSC in Exp. 1, 2, 3 and 4, respectively; n = 58, 29, 13 and 3 active HSC in Exp. 1, 2, 3 and 4, respectively; all experiments pooled: n = 24 inactive HSC and n = 103 active HSC). (**C**) Statistical comparison of LT-HSC (left) and ST-HSC (right) gene expression signatures between inactive HSC (n = 24) and active (multilineage) HSC (n = 103). (**D**) Heatmap showing the classification of inactive and active (multilineage) HSC as LT-HSC or ST-HSC based on transcriptomic data. (**E**) Statistical comparison of cell cycle scores between inactive HSC (n = 24) and active (multilineage) HSC (n = 103). (**F**) Statistical comparison of the dormant HSC signature between inactive HSC (n = 24) and active (multilineage) HSC (n = 103). (**G**) Heatmap of genes differentially expressed between active (multilineage) HSC and inactive HSC. Rows, differentially expressed genes; HSC, columns. The indicated P values were calculated using two-sided Wilcoxon–Mann–Whitney rank-sum test for **B, C, E, F**, and Fisher’s exact test for **D.** For the boxplots in **B, C, E, F**, the central line is the median, box limits are the first and third quartiles and whiskers extend to ±2.698 SD.

Next, we evaluated the cell-cycle score (table S4, Materials and Methods), which quantifies the expression of genes enriched in S, G2 and M phases of the cell cycle (*21, 22*). Inactive and active HSC did not score on average significantly different with respect to cell-cycle activity (Fig. 3E). Both active and inactive populations displayed a large spread in cell cycle score, suggesting that subsets in each population were cycling (fig. S11A). We detected on average more than one cell per clone (1.5 ±0.8 and 4.3 ±5.4, respectively, for inactive and active HSC; mean ± SD) (table S5), indicating that HSC in inactive and active clones can undergo symmetric self-renewing divisions; inactive clones, however, were significantly smaller than active ones (p = 0.02, two-sided Wilcoxon rank-sum test). Based on label retention, dormant HSC have been defined previously (*23*). Using a gene expression signature for dormant HSC (*24*) (table S4), the dormancy signature was more characteristic of active than inactive HSC (Fig. 3F), which suggests that label retaining cells may contribute to differentiation. In sum, inactive and active HSC did not fit into previously described HSC subpopulations (LT, ST or dormant).

We identified differentially expressed genes (DEG), which yielded genes more highly expressed in inactive than active HSC and vice versa (Fig. 3G; table S6). To test for robustness of differential gene expression, we performed random forest analysis which revealed separation between inactive and active HSC (with 76% probability of correct classification; fig. S12A). DEG contained previously described genes related to HSC reconstitution capacity (*Cd34*; (*25*)) and HSC maintenance under steady state (*Mycn*; (*26*)), while others were identified in stem cells of different tissues but remain poorly understood in HSC (*Bex1, Bex4*; (*27*)). Consistent with our observations on cell-cycle score and dormancy signatures, DEG were not enriched for cell-cycle regulators (and direct cell-cycle regulators which were found were expressed as a pair of activator, *Ccnd2*, and inhibitor, *Cdkn1c*, in inactive HSC) (Fig. 3G). Several DEG increased (*Cd34, Vim, Serpinb1a, Plac8*) or decreased (*Apoe, Mycn, Bex1, Pdzk1ip1, Uba7, Gng11*) their expression along pseudotime rank (fig. S11B), consistent with the enrichment of inactive HSC at the root of the transcriptional landscape (fig. S11C). The transcription factor *Hoxb2*, which is more highly expressed in inactive HSC, has previously been found to be overexpressed in acute myeloid leukemia (*28*). Therefore, Hoxb2 might be an inhibitor of HSC differentiation.

## Molecular characterization of multilineage versus restricted HSC clones

Multilineage and myelo-erythoid-restricted HSC clones realize different fates in that the former include, and the latter lack lymphoid potential (myelo-erythoid-biased HSC clones have not been further considered). To gain insights into transcriptional differences, we ranked, according to the landscape and differentiation direction shown in Fig. 2D, myelo-erythroid-restricted HSC clones relative to multilineage HSC clones, and included inactive HSC for additional reference. Restricted HSC were positioned in this pseudotime order between inactive and multilineage HSC (Fig. 4A). Restricted and multilineage HSC did not differ in cell-cycle score (Fig. 4B). Fate appeared also unrelated to clone size given that cell numbers per clone were not significantly different (4.3 ±5.4 for multilineage, and 2.5 ± 1.9 for myelo-erythoid-restricted HSC clones; table S5; p = 0.34, two-sided Wilcoxon rank-sum test). Using signatures based on differential gene expression comparing CMP and CLP (table S4, Materials and Methods), we searched for transcriptional evidence of priming towards myelo-erythroid versus lymphoid lineages at the HSC stage (Fig. 4C). The lymphoid, but not the myelo-erythroid signature, was reduced in myelo-erythroid-restricted compared to multilineage HSC (Fig. 4D). At the MPP stage, the lymphoid signature of MPP from multilineage HSC compared to MPP from myelo-erythroid-restricted HSC was more pronounced (Fig. 4E, F). We did not find differences in cell cycle score between these fate distinct MPP (Fig. 4G). Taken together, myelo-erythroid restricted clones were distinguished by low expression of the lymphoid lineage signature, which was already noticeable at the HSC stage and became clearly evident by the MPP stage. The clear distinction between myelo-erythroid and multilineage MPP was also evident by supervised classification (fig. S12B, C).

**Fig. 4.**
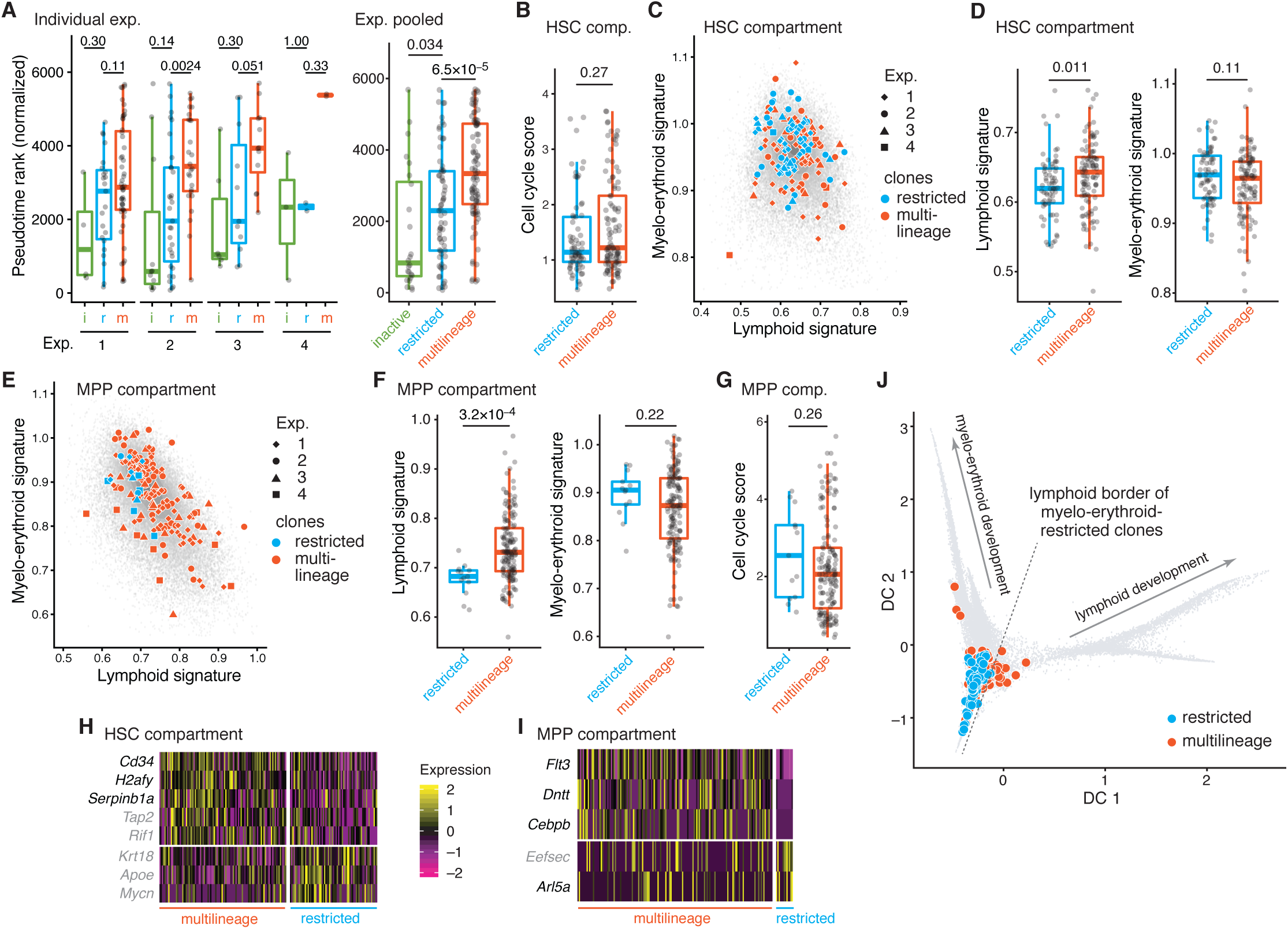
Transcriptional hallmarks of multilineage and myelo–erythroid-restricted HSC clones. **(A**) Statistical comparison of pseudotime ranks between inactive (i), myelo-erythroid-restricted (r), and multilineage (m) HSC (for individual experiments: n = 4, 11, 6 and 3 inactive HSC in Exp. 1, 2, 3 and 4, respectively; n = 25, 31, 11, 2 myelo-erythroid-restricted HSC in Exp. 1, 2, 3 and 4, respectively; and n = 58, 29, 13 and 3 active HSC in Exp. 1, 2, 3 and 4, respectively; pooled data: n = 24 inactive HSC, n = 69 myelo–erythroid-restricted HSC and n = 103 multilineage HSC). (**B**) Statistical comparison of cell-cycle scores between myelo-erythroid-restricted HSC (n = 69) and multilineage HSC (n = 103). (**C**) Scatter plot showing myelo-erythroid versus lymphoid gene signature between myelo-erythroid-restricted and multilineage HSC (colored dots, cell numbers as in (B)). The grey background distribution was computed from the scores for all single-cell transcriptomes of HSC, showing that the barcoded HSC (colored dots) are representative. (**D**) Statistical comparison of lymphoid and myelo-erythroid gene signatures between myelo-erythroid-restricted HSC (n = 69) and multilineage HSC (n = 103). **(E)** Scatter plot showing myelo-erythroid versus lymphoid gene signature of MPP derived from myelo–erythroid-restricted (n = 13) and multilineage (n = 151) HSC clones. The grey background distribution was computed from the scores for all single-cell transcriptomes of MPP, showing that the barcoded MPP (colored dots) are representative. (**F**) Statistical comparison of lymphoid and myelo–erythroid gene signature of MPP derived from myelo-erythroid-restricted and multilineage HSC clones (cell numbers as in (E)). **(G)** Statistical comparison of cell-cycle scores between MPP that are derived from myelo-erythroid-restricted and multilineage HSC clones (cell numbers as in (E)). **(H)** Heatmap of genes differentially expressed between myelo-erythroid-restricted HSC (n = 69) and multilineage HSC (n = 103). (**I**) Heatmap of genes differentially expressed between MPP derived from myelo-erythroid-restricted (n = 13) and multilineage (n = 151) HSC clones. (**J**) Projection of HSC and MPP from myelo-erythroid-restricted (blue) or multilineage (red) HSC clones onto the transcriptional landscape. Data shown are from Exp 1, 2, and 3. P values were calculated using two-sided Wilcoxon–Mann–Whitney rank-sum test for (A), (B), (D), (F), (G). For the boxplots in (A), (B), (D), (F), (G), the central line is the median, box limits are the first and third quartiles and whiskers extend to ±2.698 SD. In (H) and (I), genes in black satisfy FDR < 5% and genes in grey satisfy FDR < 20%.

Differentially expressed genes between multilineage and myelo-erythroid-restricted HSC clones included transcripts more abundant in multilineage HSC, and vice versa (Fig. 4H, table S6). High *Cd34* and *Serpinb1a* expression as well as low *Mycn* and *Apoe* expression distinguished active multilineage HSC from both inactive (Fig. 3G, table S6) and myelo-erythroid-restricted HSC (Fig. 4H, table S6). By contrast, differential expression of *H2afy, Tap2, Rif1* and *Krt18* was specific to the distinction between multilineage and myelo-erythroid restricted HSC. The *H2afy* gene, coding for a histone H2 variant, has previously been implicated in HSC maintenance and balanced output (*29, 30*). Genes differentially upregulated in MPP from multilineage HSC clones included lymphoid marker genes *Flt3* and *Dntt* (Fig. 4I, table S6) expressed in lymphoid-biased MPP (LMPP) (*31*), indicating that phenotypic LMPP arise from multilineage HSC clones. Consistent with this finding, we observed a boundary in the transcriptional landscape between myelo-erythroid and lymphoid development that was crossed in the MPP region only by cells of multilineage clones (Fig. 4J).

## Discussion

HSC and progenitor fates have traditionally been studied by phenotype-driven prospective isolation, which included fate readout by transplantation (*32*). Correlations between transcriptomes and fates, determined by transplantation or in vitro assays, have recently been studied in split-daughter-cell experiments (*33*). The development of *PolyloxExpress* now allows linking transcriptomes and fates under physiological conditions. Applying *PolyloxExpress* to hematopoiesis, we find that the adult hematopoietic system retains a functional imprint of its clonal development in the fetus. Barcoding and fate mapping revealed consistent fates of developmentally defined HSC clones (differentiation-inactive, multilineage, myelo-erythroid-restricted), and HSC clones with common fate were distinct on the transcriptional landscape. While multilineage clones could be composed of cells with mixed fates, analysis of variance showed homogeneity of transcriptomes within each fate category, including multilineage, which provides support for clonal coherence. While a single time point of analysis reveals the cumulative output from HSC, analyzing pathways from HSC via fast turning over progenitors to peripheral lineages offers in addition the view on fates currently realized by HSC clones in adult bone marrow. The fact that progenitor generation with no exception fit the peripheral lineage supports stability of clonal HSC fates, i.e., inactive HSC clones did not only lack mature cells but also progenitors, and myelo-erythroid-restricted HSC clones contained myeloid but lacked lymphoid progenitors. In further support, myelo-erythroid-restricted HSC clones have not produced lymphocytes, and inactive HSC clones none of the sampled lineages, between the time of label induction in the embryo until the time of analysis (between 7 and 20 weeks after birth). As a caveat, inactive HSC clones may have generated lineages not sampled here, such as mast cells, innate lymphocytes, or megakaryocytes. Given that inactive HSC also did not produce MPP, such lineages would need to be generated independent of MPP. Fate stability indicates coherent, or stereotypical behavior of cells in a clone, which was also previously suggested from color barcoding and tracing experiments (*34*). Stable fate biases of HSC have previously been described following transplantation (*10, 35-39*), and have included, besides multilineage and myelo-erythroid-biased HSC, lymphoid-biased HSC. As previously shown for mice around one year of age (*12*); we did not find lymphoid-biased HSC clones up to 20 weeks of age, indicating that lymphoid fate bias may be more pronounced after transplantation (*40*).

*PolyloxExpress* may be generally applicable to probing the transcriptome of fate-defined stem cells which continuously support tissue renewal, i.e. in organs in which stem cells and their products are available for parallel analysis. Towards this aim, CRISPR/Cas9-based barcoding approaches, hitherto used for developmental lineage reconstruction (*41-43*), might also be adaptable (*44*).

Barcode analysis combined with single cell RNA sequencing provided transcriptome information of HSC with known fates. This enabled positioning of functionally different HSC on the transcriptional landscape, with inactive, active myelo-erythroid restricted and active multilineage HSC residing at partially overlapping but overall distinct locations. Although pseudo-time rank increased comparing inactive to active multilineage HSC clones, our results suggest that clone location in the transcriptional landscape may be stable, rather than clones migrating along developmental progression. In particular, the fraction of inactive clones did not decrease with age of the mice, being 15±8% (mean ± SD) for mice of 7-20 weeks of age studied here, and 20% and 22% for mice of ages 38 and 47 weeks, respectively (*12*). Also, the fraction of myelo-erythroid-restricted clones did not decrease with age. Differentiation-inactive HSC clones are not proliferation-inactive, and a reported dormancy HSC signature (*24*) was more characteristic of differentiation-active than inactive HSC, suggesting that clones of the three types of fate studied here may each contain label-retaining HSC. Transcriptional differences detected between HSC clones with distinct fates may aid future studies into the molecular regulation of stem cell activity.

## Supporting information

Supplement Pei et al.

## ACKNOWLEDGEMENTS

We thank Frank van der Hoeven and Ulrich Kloz (DKFZ Transgenic Service Facility) for help with gene targeting, the DKFZ Genomics and Proteomics Core Facility for providing Illumina sequencing services, Jan-Philipp Mallm, Katharina Bauer, and Michelle Liberio (DKFZ Single-cell OpenLab) for discussions and technical support, Jens Rössler for discussions, Günter Küblbeck, Nina Claudino and Sven Schäfer (DKFZ) for expert technical assistance, and all members of the Rodewald laboratory for ongoing support and discussions.

## Author contributions

W.P. and F.S. performed most experiments, X.W. performed most computational analysis with support from A.G. and T.H., A.K.F., K.B., K.K., Q.Z. made experiments, C.Q. performed S.M.R.T. sequencing, S.S. supported the project from the Scientific Genomics Platform a the BIMSB/BIH, T.B.F. designed the PolyloxExpress cassette and made experiments, T.B.F., T.H. and H.R.R. supervised the study.

## Author information

The authors declare no competing financial interests.

## Data and material availability

All data, codes, and materials used in this study are available. Please contact H.R.R and T.H. for material requests.

## SUPPLEMENTARY MATERIALS

Materials and Methods

Supplemental Figure Legends

Supplemental Figures (Fig. S1 to S12)

Supplemental Tables (Tables S1 to S6)

