## Supplement Pei et al. for "Resolving fate and transcriptome of hematopoietic stem cell clones"

#### Materials and Methods

**Mice.** *Rosa26*$^{PolyloxExpress}$ (*B6-Gt(ROSA)26Sor*$^{tm2.1(PolyloxExpress)Hrr}$), *Tie2*$^{MeriCreMer}$ (*Tek*$^{tm1.1(icre/Esr1*)Hrr}$) (*1*), C57BL/6 and CD1 strains were used in this study. *Rosa26*$^{PolyloxExpress}$ knock-in mice were generated by conventional gene targeting as described below. All mice were kept in individually ventilated cages under specific pathogen-free conditions in the animal facility at the German Cancer Research Center (DKFZ, Heidelberg). Male and female mice were used, no randomization was done, no blinding was done and no animals were excluded from the analysis. No statistical methods were used to predetermine sample size. All animal experiments were performed in accordance with institutional and governmental regulations, and were approved by the Regierungspräsidium (Karlsruhe, Germany).

**Generation of *Rosa26*$^{PolyloxExpress}$ knock-in mice.** The *PolyloxExpress* cassette was designed on the basis of the *Polylox* cassette (*2*); details are shown in fig. S1A. In brief, the *PolyloxExpress* cassette consists of a tandem-dimer tomato (tdTomato) fluorescent reporter followed by a 2A peptide and two translational stop codons, the *Polylox* sequence (identical to the original *Polylox* sequence except for a 64-bp truncation at the 5’ end), and a bovine growth hormone polyadenylation (pA) signal. This cassette was introduced into the *Rosa26* locus in murine embryonic stem (ES) cells by conventional gene targeting via homologous recombination. The targeting vector consists of a homologous short-arm, a neomycin resistance gene (*Neo*) flanked by FRT sites, the *PolyloxExpress* cassette as described above, a homologous long-arm, and the diphtheria toxin subunit A (DT-A) driven by the *PGK1* promoter for selection against random integration. The targeting vector was electroporated into ES cells. Transfected ES cell clones were selected by G418, and screened by PCR using the Expand Long Template PCR System (Sigma) with PCR buffer supplemented with 4% DMSO and the primers HL16 (5′-CCTAAAGAAGAGGCTGTGCTTTGG-3′) and HL15 (5′-AAGACCGCGAAGAGTTTGTCC-3′)(*3*). PCR-positive clones were validated by Southern blot using a Dig-labeled 820 bp external probe (*3*). To enable tdTomato and *PolyloxExpress* expression, the *Neo* cassette was removed *in vitro* from *Rosa26*$^{Neo\text{-}PolyloxExpress}$ ES cells by transient *Flp* recombinase transfection. ES cell clones were tested for *Neo* deletion by PCR using the primers #2,656 (5’-AGGGAGCTGCAGTGGAGTAG-3’) and #2,657 (5’-ACCATGGTCTTGGAGAAACAG-3’).

*Neo*-deleted *Rosa26*$^{PolyloxExpress}$ clones were injected into C57BL/6 blastocysts and chimeric male offspring were backcrossed to C57BL/6 mice for germline transmission of the *Rosa26*$^{PolyloxExpress}$ allele. Genotyping of *Rosa26*$^{PolyloxExpress}$ mice is done by PCR using the primers #1,116 (common FWD primer) 5'-AAGGGAGCTGCAGTGGAGTA-3', #1,117 (wt Rev primer) 5'-TAAGCCTGCCCAGAAGACTCC-3' and #1,118 (KI Rev primer) 5'-AAGACCGCGAAGAGTTTGTCC-3', yielding 210 bp wild type or 300 bp knock-in products. Both heterozygous *Rosa26*$^{PolyloxExpress/+}$ and homozygous *Rosa26*$^{PolyloxExpress/PolyloxExpress}$ mice are viable and fertile, and do not show any obvious phenotype.

**In vitro recombination of *PolyloxExpress* barcodes in ES cells**. For the *in vitro* recombination of *PolyloxExpress* barcodes, MeriCreMer-transfected *Rosa26*$^{PolyloxExpress}$ ES cell clones were generated. Therefore, *Rosa26*$^{PolyloxExpress}$ ES cells were transfected with a pAN-MiCM-neomycin expression vector, carrying the tamoxifen-inducible MeriCreMer (*4*) under the control of the human *ACTB* promoter. Transfected ES cells were selected with G418 and screened for stable *MeriCreMer* integration by PCR using forward (5'-AAGAACCTGATGGACATGTTCAGG-3') and reverse (5'-TCTGTCAGAGTTCTCCATCAGGGA-3') primers yielding 390 bp product. For the induction of barcode recombination, cells were incubated with 4-hydroxy-tamoxifen (4-OHT) for the indicated times. In the bulk analysis cell aliquots were harvested directly after induction for DNA or RNA extraction (fig. S2A and B). For clonal barcode read out, ES cells were washed after 4-OHT induction and chased for 16 days before individual ES cell clones were picked for further expansion and DNA or RNA extraction (fig. S2C and D). Genomic DNA was extracted by using phenol-chloroform. RNA was purified by RNeasy Plus Mini Kit (Qiagen) and used for cDNA synthesis with oligo(dT) (Invitrogen). The *PolyloxExpress* cassette was amplified by locus-specific PCR with the Expand Long Template PCR system using 200-300 ng genomic DNA template, or 20-50 ng cDNA template and the following conditions: 5 min at 95 °C; (30 s at 95 °C, 30 s at 54 °C, 3 min at 72 °C) 35 times; 10 min at 72 °C with primers #2,653 (5'-CGAACTTCTCTCTGTTAAAGCAAG-3') and #2,427 (5'-CATACCTTAGAGAAAGCCTGTCGAG-3'). RT-PCR conditions for the amplification of *Actb* were: 5 min at 95 °C; (30 s at 95 °C, 30 s at 57 °C, 30 s at 72 °C) 35 times; 10 min at 72 °C with primers #2,609 (5'-ACGCAGCTCAGTAACAGTCC-3') and #2,610 (5'-CATCACTATTGGCAACGAGC-3'). PCR products were analyzed by gel electrophoresis and Sanger sequencing (GATC Biotech).

**Barcode induction in mice.** For barcode induction in HSC precursors at E9.5 of embryonic development, timed matings were set up between *Rosa26*$^{PolyloxExpress/PolyloxExpress}$ and *Tie2*$^{MeriCreMer/+}$ mice. Nine days after the day of the plug (day 0.5), pregnant mice were treated by oral gavage with a single dose of 2.5 mg tamoxifen and 1.25 mg progesterone in peanut oil (all from Sigma). The pups were delivered on E19.5 by caesarean section, raised by CD1 foster mothers, and genotyped for the *Tie2*$^{MeriCreMer}$ and *Rosa26*$^{PolyloxExpress}$ alleles, 4 weeks after birth as described (*5*).

**Fluorescence-activated cell sorting (FACS).** The following antibodies were used: CD3ε-PerCP-Cy5.5 (145-2C11), CD4-BV421 (GK1.5), CD8a-BV421 (53-6.7), CD11b-BV421 (M1/70), CD19-BV421 (6D5), CD44-BV605 (IM7), CD71-PE-Cy7 (RI7217), CD115-BV605 (AFS98), CD117-BV711 (2B8), CD150-PE-Cy7 (TC15-12F12.2), CD150-BV605 (TC15-12F12.2), Gr1-BV421 (RB6-8C5), NK1.1-BV421 (PK136), Ter119-BV421 (Ter119) from Biolegend. BP1-FITC (6C3), CD4-PerCP-Cy5.5 (RM4-5), CD4-APC (RM4-5), CD11b-PE-Cy7 (M1/70), CD24-APC-eFluor780 (M1/69), CD34-eFluor660 (RAM34), CD45-AlexaFluor700 (30-F11), CD45R-PerCP-Cy5.5 (RA3-6B2), CD48-FITC (HM 48-1), CD127-PE-Cy7 (A7R34), CD135-APC (A2F10), MHCII-APC (M5/114.152), Sca1-PerCP-Cy5.5 (D7) from Life Technologies. CD3ε-FITC (17A2), CD8a-FITC (53-6.7), CD11b-FITC (M1/70), CD16/32-BV605 (2.4G2), CD19-FITC (1D3), CD19-APC (1D3), CD25-PE-Cy7 (PC61), CD43-APC (S7), Gr1-FITC (RB6-8C5), Ly6C-APC-Cy7 (AL-21), Ly6G-PerCP-Cy5.5 (1A8), NK1.1-FITC (PK136) from BD Pharmingen.

Bone marrow cells were harvested from femora, tibiae, fibulae, pelvis, humeri, radiuses, ulnae and spine by crushing the bones in FACS staining buffer (5% fetal bovine serum in DPBS) with mortar and pestle and filtering through a 40 μm mesh (BD Falcon). Spleen and thymus were dissociated by passing the cells through a 40 μm filter with FACS staining buffer. Cell suspensions were blocked with 300 μg/ml whole mouse IgG (Jackson ImmunoResearch Laboratories) for 15 min, followed by 45 min staining with concentration titrated fluorescent-conjugated antibodies on ice.

Before FACS analysis, Sytox Blue (Invitrogen) solution (1:10.000) was added for dead cell discrimination. Lineage depletion for the enrichment of HSC and progenitors was conducted before fluorescent antibody staining and cell sorting as previously described (*5*).

All cells were purified by cell sorting on a BD FACS Aria III flow cytometer (fig. S5). Cell populations were defined by combination of the following markers: HSC (Lin$^-$ Sca1$^+$ Kit$^+$ CD48$^-$ CD150$^{+/-}$), MPP (Lin$^-$ Sca1$^+$ Kit$^+$ CD48$^+$ CD150$^-$), CMP (Lin$^-$ Sca1$^-$ Kit$^+$ CD16/32$^{med}$ CD34$^{med}$), GMP (Lin$^-$ Sca1$^-$ Kit$^+$ CD16/32$^+$ CD34$^+$), MEP (Lin$^-$ Sca1$^-$ Kit$^+$ CD16/32$^-$ CD34$^-$), CLP (Lin$^-$ Kit$^{lo}$ Sca1$^{lo}$ CD127$^+$ CD135$^+$), pro-B cells (combined fractions B and C) (CD4$^-$ CD8$^-$ CD11b$^-$ Ter119$^-$ Gr-1$^-$ Nk1.1$^-$ CD43$^{med}$ CD45R$^{med}$ CD24$^+$ BP1$^-$ and BP1$^+$), granulocytes (Gr) (CD4$^-$ CD8$^-$ CD19$^-$ CD11b$^+$ Gr-1$^+$), monocytes (CD4$^-$ CD8$^-$ CD19$^-$ Ter119$^-$ CD45$^+$ MHCII$^-$ Ly6G$^-$ Ly6C$^+$ CD11b$^+$ CD115$^+$ ), CD4+ T cells (CD4$^+$ CD8$^-$ CD19$^-$ CD11b$^-$ Gr-1$^-$), CD8+ T cells (CD4$^-$ CD8$^+$ CD19$^-$ CD11b$^-$ Gr-1$^-$), B cells (CD4$^-$ CD8$^-$ CD19$^+$CD11b$^-$ Gr-1$^-$), erythroid progenitor (EryP) stage II (basophilic erythroblasts: CD3ε$^-$ CD11b$^-$ CD19$^-$ Gr1$^-$ NK1.1$^-$ Ter119$^+$ CD71$^+$ CD44$^+$ FSC$^+$) and stage III (polychromatic erythroblasts: CD3ε$^-$ CD11b$^-$ CD19$^-$ Gr1$^-$ NK1.1$^-$ Ter119$^+$ CD71$^+$ CD44$^{med}$ FSC$^{med}$), pre-T cells as (CD3ε CD4 CD8 CD11b CD19 Ter119 Gr1 NK1.1)-negative CD25$^+$ CD44$^+$ DN2 and CD25$^+$ CD44$^-$ DN3 fractions.

**Detection of DNA barcodes in bulk samples by single molecule real-time (SMRT) sequencing.** Genomic DNA was prepared from sorted cells (30,000 cells) by proteinase K treatment and heat inactivation as described (*5*). The *PolyloxExpress* cassette was amplified by PCR as described above for the in vitro recombination in ES cells. PCR products were cleaned up with 0.7 x AMPure beads (Beckman Coulter) and eluted in 25 µl EB buffer. SMRTbell library preparation, SMRT sequencing and generation of CCS reads were performed as described in (*5*).

**RNA barcode and transcriptome detection in single cells.** Live cells (9,000 – 20,000 cells) were purified by cell sorting, and single cells were captured using the Chromium System according to the Chromium Single Cell 3’ Reagent Kit v3 protocols (10x Genomics). In brief, single cells were encapsulated into individual droplets together with indexed beads. In each droplet, mRNA was released from a single cell, and captured by a bead containing oligo anchors composed of oligo(dT), cell-specific index, Unique Molecular Identifiers (UMI) and a PCR adaptor sequence. After reverse transcription in the droplets, cDNA was released, cleaned up according to the Chromium Single Cell 3' Reagent Kits protocol, and amplified using read 1 and read 2 primers (10x Genomics). The amplified 10x cDNA library was used for RNA barcodes retrieval and transcriptome library preparation.

**Pre-processing of single-cell RNA-seq data.** In total, in each of the four experiments, we generated 2.37, 3.87, 2.92, and 2.18 billion paired-end Illumina sequencing reads from the prepared single-cell RNA-seq (scRNA-seq) libraries. Sequencing reads from individual samples were first subjected to Cell Ranger (version 3.1.0; 10x Genomics) pipeline analysis. In brief, the read parts derived from RNA molecules were aligned to the mouse reference genome (mm10) using splicing-aware aligner STAR (version 2.5.1b) (*6*); the read parts containing the information on cell indexes and unique molecular identifiers (UMI) were used, respectively, to distinguish cells and to count RNA molecules. The distribution of UMI counts per cell index displayed bimodality, so that the clearly recognizable valley was used to separate authentic cells and background cell indexes. As a result, Cell Ranger recovered 37,772, 78,651, 48,028 and 42,225 cells in each of the four experiments. With the resulting gene-by-cell UMI count matrixes, we performed basic data analysis using the *Seurat* R package (version 3.1.1) (*7*). In short, we filtered out low-quality cells and possible cell doublets as cells not satisfying the following criteria: (1) number of detected genes between 1800 and 8000, (2) number of total UMIs less than 80,000, and (3) percentage of UMIs derived from mitochondrial genes below 8%. This step discarded on average approximately 8% of cells recovered by Cell Ranger, most of which were due to too few detected genes. We then normalized the UMI counts by library size factors, where the library size factor is equal to the sum of UMIs in each single cell divided by 10,000. Subsequently, we log2-transformed the normalized UMI counts (plus one pseudocount), such that the final transformed expression values were $e_{ij} = \log2(\frac{C_{ij}}{\sum_i C_{ij}} \times 10000 + 1)$, where $C_{ij}$ denotes the UMI count of gene $i$ in cell $j$.

To mitigate against unwanted technical variation in the scRNA-seq data, we adopted the standard procedure of selecting 1000 highly variable genes (HVGs) that were informative of the biological variability in the transcriptional landscape. We adopted the variance-stabilizing transformation (vst)-based method implemented in the *Seurat* v3 package for this purpose (*8*). In brief, variance-stabilizing transformation of raw count data was applied to account for the inherent mean-variance relationship in scRNA-seq, and the variance of standardized values across all cells was computed for each gene. We then ranked the standardized variance, and the top 1000 genes were selected for the reconstruction of hematopoietic transcriptional landscape.

**Retrieval of single-cell *PolyloxExpress* barcodes.** Single-cell *PolyloxExpress* barcodes were determined from RNA by SMRT sequencing of the *Polylox* cassette together with cell indexes (CIs; each a 16 bp stretch), introduced via indexed beads in individual encapsulated droplets (10x Genomics). Taking circular consensus sequencing (CCS) reads from SMRT sequencing, we first filtered for the expected segmental structure, which includes the 5′-end of the *Polylox* cassette, the *Polylox* cassette itself and 3′ end (in this order), followed by a stretch of adenines (As) and a primer sequence to identify CIs (fig. S3C). Only CCS reads of correct structure were used for further processing. We recorded barcodes retrieved from each read using the RPBPBR pipeline (*5*) together with CIs. After an additional filtering step to remove a small number of CIs unobserved in the whole-transcriptome sequencing data and illegitimate *PolyloxExpress* barcodes, we aggregated *PolyloxExpress* barcodes by CIs.

In some CI aggregates, we observed multiple *PolyloxExpress* barcodes, partially because of sequencing errors in CIs, random association of CIs and *PolyloxExpress* barcodes due to PCR chimeras, or doublets captured by the 10x Genomics platform. To identify a unique barcode per CI, we considered only the most abundant barcodes that satisfied the following two criteria: (1) the abundance level is sufficient (supported by >80% reads in the CI aggregate), and (2) the association of *PolyloxExpress* barcodes and CIs is not observed by chance.

**Rare *PolyloxExpress* barcodes.** Barcode generation probabilities (*Pgen*) have been described for *Polylox* barcodes (*2*), and apply for *PolyloxExpress* barcodes. Barcodes with a *Pgen* lower than $5 \times 10^{-4}$ were regarded as rare barcodes, generated with probabilities > 91% in only a single precursor cell (*5*).

**Fate classification of HSC clones by barcode propagation.** In each of the four experiments, barcodes with $P_{\text{gen}} < 5 \times 10^{-4}$ and detected in HSC or MPP were compared in HSC, progenitors, myeloid, erythroid, and lymphoid lineages (all populations indicated in Fig. 2A and fig. S7C, D).

For each cell population, barcode counts obtained from sample repeats were combined, and barcode frequencies were calculated as barcode counts divided by the total counts (without barcode filtering). Barcode frequencies are displayed as heatmaps (Fig. 2A and fig. S7C, D). Based on the presence or absence of barcodes in peripheral lineages (Gr, Mono, EryPII/III, CD4, CD8, and B), HSC clones were sorted into inactive, myelo-erythroid-restricted, myelo-erythroid biased and multilineage fates. Barcodes with $P_{\text{gen}} > 5 \times 10^{-4}$ showing no output (inactive) or myelo-erythroid-restricted output are included, and shown in red font, while the barcodes with $P_{\text{gen}} < 5 \times 10^{-4}$ are shown in black font. To gain an overview of the peripheral barcode distribution, barcodes with $P_{\text{gen}} < 5 \times 10^{-4}$ and detected in two independent sample replicates were compared between peripheral lineages (populations as indicated in fig. S7A). Barcode frequencies were calculated in the same way described above and are displayed as heatmaps (fig. S7A). Hierarchical clustering was performed based on barcode rank correlations between peripheral lineages (fig. S7B), as described previously (*2*).

**Reconstruction of the transcriptional landscape using the diffusion map.** Cells at the same cell-cycle stages tend to have similar transcriptome profiles dominated by cell-cycle-related genes, which may confound the ordering of cells in the transcriptional landscape according to developmental transitions. To avoid this effect, we first adjusted gene expression profiles by linearly regressing out cell-cycle scores (S scores and G2M scores; see below) (*7, 9*). Let $s_j$ and $m_j$ denote the S score and G2M score of cell $j$, respectively, then the normalized expression $e_{ij}$ of gene *i* can be modelled as $e_{ij} = \alpha_i \cdot s_j + \beta_i \cdot m_j + \varepsilon_{ij}$, where $\alpha_i$ and $\beta_i$ are regression coefficients. By minimizing $\sum_j {\varepsilon_{ij}}^2$, the coefficients can be resolved and gene expression predicted by cell cycle scores can be obtained $\hat{e}_{ij} = \hat{\alpha}_i \cdot s_j + \hat{\beta}_i \cdot m_j$. To remove the impact of cell cycle on gene expression, the estimated residuals $\hat{\varepsilon}_{ij} = e_{ij} - \hat{e}_{ij}$ were used for the transcriptional landscape analysis.

To reconstruct the transcriptional landscape, we used the diffusion map, which is a nonlinear dimensionality reduction approach focusing on learning the underlying manifold from observed data (*10*). We used the implementation in the *destiny* R package (v3.0.0) to build the transition matrix that describes random walks between cells and calculate the dominant eigenvectors for visualizing the underlying embedding in low-dimensional space. Cell-cycle-adjusted and Z-score-normalized expression values of 1000 HVGs were used to compute the diffusion maps. For visualization, the data were plotted in a coordinate system spanned by the first two diffusion components (ordered by eigenvalues in descending order).

**Diffusion pseudotime analysis**. We used diffusion pseudotime (DPT) to order the cells along putative differentiation trajectories and identify branch points of these trajectories (*11*). Notably, this algorithm works on the full dimensions of the diffusion-map space, rather than the low-dimensional embedding used for visualizing the transcriptional landscape, allowing the detection of subtle changes in gene expression during development. In this study, we adopted the implementation of DPT algorithm in the *destiny* R package (v3.0.0) to compute DPT for each cell with respect to a specified root cell, a tip HSC, and the DPT was used to quantify developmental progression in hematopoiesis.

To identify branch points, we relied on the criterion of switching DPT correlations (*11*). We applied this criterion iteratively, first identifying the major lymphoid and myelo-erythroid branches, and then downstream sub-branches. All the identified branches were then annotated by gene expression of lineage markers (table S3).

**Calculation of cell-cycle scores.** To quantify cell-cycle status in our data, we adopted the S- and G2/M-phase gene sets used in previous studies (*12*), which have been shown to characterize well S or G2/M phases in both synchronized cell-population (*13*) and single-cell (*14*) experiments. The genes used to calculate the cell-cycle scores are listed in table S4. We adapted the framework of gene set enrichment analysis (GSEA) (*15*) to quantify cell cycling status for each cell: We first calculated the enrichment scores and then normalized by the mean of enrichment scores computed based on permuted gene-by-cell matrices. The normalized enrichment scores were taken as cell-cycle scores.

**Calculation of phenotypic gene signatures.** Phenotype-associated gene signatures are useful to characterize single-cell transcriptomes. The gene lists used to calculate LT-HSC and ST-HSC signatures (table S4) were extracted by comparing gene expression patterns between phenotypically sorted LT-HSC and ST-HSC profiled by scRNA-seq (Fanti et al. unpublished). Normalized expression values of individual genes were tested for differential expression between the two cell populations using Wilcoxon rank sum tests, and the raw *P*-values were corrected for multiple tests using the Benjamini-Hochberg procedure. Genes with >1.2-fold difference between populations and adjusted *P*-value < 0.05 were used for gene signature calculation. Similarly, the myelo-erythroid and lymphoid signatures were extracted by comparing gene expression profiles between phenotypically sorted CMP and CLP. As the gene expression differences between CMP and CLP were more prominent, we applied a more stringent threshold (> 1.5 fold) with the same adjusted *P*-value threshold (< 0.05). The dormant HSC signature was extracted from differential gene expression analysis published previously (*16*). We took genes with adjusted *P*-value < 0.01 and log2(dHSC/aHSC) > 2 as dormant HSC genes, where dormant HSC (dHSC) and active HSC (aHSC) were defined in the original paper via retention of proliferation-dependent cell label. For all gene sets, we calculated the gene signatures as the average normalized expression $\frac{1}{n}\sum_{i\in G} e_{ij}$, where $e_{ij}$ is the normalized expression value (defined above), *G* denotes a gene set of interest, and *n* is the size of this gene set. All genes used for gene signature calculation in this study are listed in table S4.

**Prediction of LT-HSC and ST-HSC.** Our sorted, fate- and transcriptome-analyzed HSC (clonal data set) contained both phenotypic long-term (LT) and short-term (ST) cells. To assign single HSC in this data set by transcriptome to LT or ST categories, the *SingleR* approach **(*17*)** was used. As a reference, we employed separate scRNA-seq data from sorted LT-HSC and ST-HSC (reference data set; Fanti et al. unpublished). To overcome dropout issues of scRNA-seq data as transcriptome reference, we aggregated 500 random single cells of each reference population to create 100 pseudo-bulk samples. Then, each HSC transcriptome of clonal data set was compared to 100 LT and 100 ST pseudo-bulk transcriptomes via Spearman correlation coefficients, and the 80th percentile of the coefficients per cell type was considered for cell type assignment.

**Identification of differentially expressed genes.** To elucidate transcriptome differences between cells of different clonal fates, we adopted a widely-used approach provided in the *limma* R package (v3.42.0) (*18*) to identify differentially expressed genes (DEGs) from the whole transcriptome. The algorithm builds gene–wise linear models which consider the variables of interest as well as covariates in complete experiments. Thus, when dealing with the pooled data from four experiments in this study, we took mouse identifiers as a covariate, and tested expression difference for each gene with respect to clonal fates (including active vs. inactive, multilineage vs. restricted).

The test statistics were then subjected to an empirical Bayes procedures for information borrowing to improve statistical power and accuracy. Finally, raw $P$-values were adjusted for FDR using the Benjamini-Hochberg approach. We applied a stringent criterion for DEGs with FDR < 5% (shown in black in Fig. 3G, Fig. 4H and I) and a more relaxed criterion with FDR < 20% (shown in grey in Fig. 4H and I). All DEGs are listed in table S6.

**Fate classification of HSC and MPP by transcriptome.** To examine whether single HSC can be assigned by their transcriptomes to inactive or active (multilineage) clones, and whether single HSC or MPP to multilineage or myelo-erythroid-restricted clones, we evaluated the accuracy of cell classification based on transcriptome data. We used the random forest algorithm (implemented in *randomForest* R package v4.6-14), a supervised classification model, to build classifiers using all but one cell, which were then used to classify the remaining cell. After repeating this procedure for every HSC or MPP derived from myelo-erythroid-restricted and multilineage clones, we compared the classification outcomes and their original class labels for classification performance evaluated using receiver operating characteristic curves.

**Changes in gene expression along diffusion-pseudotime rank.** Given that inactive and active HSC have distinct positions on the transcriptional landscape (i.e. distinct pseudotime ranks), it is informative to identify genes exhibiting expression changes along the pseudotime axis. For this purpose, we regressed the expression value of each gene on the pseudotime rank using a general additive model (GAM) implemented in the *gam* R package (v1.16.1), which allows for non-linear gene expression changes. The 30 most significant genes are shown in fig. S11B and C.

#### Supplementary Figure Legends

**Fig. S1. Generation of the *Rosa26*$^{\mathit{PolyloxExpress}}$ locus**. (**A**) Gene targeting of the *Rosa26*$^{\mathit{PolyloxExpress}}$ locus. The *PolyloxExpress* cassette was inserted into the *Rosa26* locus in mouse embryonic stem (ES) cells by homologous recombination. Shown are, from top to bottom, the wild-type *Rosa26* locus, the targeting vector and targeted *Rosa26*$^{\mathit{Neo\text{-}PolyloxExpress}}$ locus. Neomycin resistance gene (*Neo*), used for the selection of targeted ESC clones, was deleted (*ΔNeo*) to generate the *Rosa26*$^{\mathit{PolyloxExpress}}$ locus. SA, short-arm; LA, long-arm; FRT, FLP recombinase target; DT-A, diphtheria toxin A. (**B**) Southern blot of wild type (WT), knock-in (KI) and three Neomycin-deleted KI (*ΔNeo*-KI) on genomic DNA from mouse tails. Gene targeting and *Neo* deletion were confirmed by the size of restriction fragments corresponding to WT (5,855 bp), KI (4,050 bp), and *ΔNeo*-KI (10,008 bp) loci using the probe indicated in (A). (**C**) Tomato expression in $CD8^+$ T cells, B cells and granulocytes measured by flow cytometry. Positive signals in PE channel indicate transcription from the *Rosa26*$^{\mathit{PolyloxExpress}}$ locus.

**Fig. S2. DNA and RNA barcode induction in vitro and demonstration of single barcode per cell**. (**A**) Experimental workflow for testing barcode recombination in *MeriCreMer*-transfected *Rosa26*$^{\mathit{PolyloxExpress}}$ ES cells. ES cells were treated for the indicated time with 4-hydroxy-tamoxifen (4-OHT). Bulk DNA and RNA were purified after the indicated periods of treatment. (**B**) Kinetics of barcode recombination in bulk DNA and RNA samples (shown in A) analyzed by gel electrophoresis. RT, reverse transcription; EtOH, vehicle control. (**C**) Comparison of DNA and RNA barcodes in ES cell clones. Induced single-cell derived ES cell clones were picked after the indicated pulse-chase treatment. After clonal expansion, DNA and RNA from the same clone were purified for *PolyloxExpress*-specific PCR amplification, followed by gel electrophoresis and Sanger sequencing analysis. (**D**) Gel electrophoresis showing identical bands for DNA and RNA barcodes in each of the 39 ES cell clones from (C).

**Fig. S3. Integrating *Polylox* barcodes with transcriptomes in single cells.** (**A**) Barcode induction in *MeriCreMer*-transfected *Rosa26*$^{\mathit{PolyloxExpress}}$ ES cells, and integrative single-cell barcode and transcriptome analyses. Cell-indexed cDNA libraries prepared on the 10x Genomics platform were split for both transcriptome analysis (Illumina) and *Polylox* barcode analysis (SMRT sequencing). (**B**) Venn diagram showing cell index (CI) consistency between whole transcriptome analysis (Illumina) and *Polylox* barcode analysis (SMRT sequencing). (**C**) Computational workflow for *Polylox* barcode retrieval and integration with transcriptome data. Step 1, filtering PacBio CCS reads for correct sequence structure of *Polylox* barcodes indexed on the 10x Genomics platform. Truncated and chimeric reads (ca. 6% of total PacBio CCS reads) as well as mouse transcriptome reads (ca. 65% reads) were discarded. After *Polylox* barcode retrieval using the RPBPBR pipeline, *Polylox* barcodes associated with the same CI were aggregated. As a result, approximately 25% of the CIs detected in Illumina sequencing were retained. Step 2, identification of unique *Polylox* barcodes in each CI aggregate. If more than one barcode was observed in a CI aggregate, the most abundant *Polylox* barcode (occurring > 80%) was assumed to be the authentic barcode, which was subjected to further statistical tests to exclude observing the CI-assigned barcode simply by chance (FDR < 1%). The CIs without identified *Polylox* barcodes were kept only for transcriptomic analysis (ca. 2%), and the remaining ca. 23% CIs were used for integrative analysis.

**Fig. S4. Barcode induction controls in vivo**. CD4 T cells, B cells and granulocytes were harvested from three different mice for barcode recombinations. The $Tie2^{MCM}Rosa26^{PolyloxExpress}$ mouse without tamoxifen (TAM) treatment (left three lanes), and the $Rosa26^{PolyloxExpress}$ mouse (middle three lanes) showed no spontaneous recombination, while the embryonically induced $Tie2^{MCM}Rosa26^{PolyloxExpress}$ mouse (right three) showed recombined DNA fragments.

**Fig. S5. Overview of FACS gating strategies**. (**A** to **I**). Surface marker combinations used for isolation of indicated cell populations. Surface markers indicated above the first plot of each panel were used to pre-define the indicated population. Pre-gatings for size (FSC, SSC) and live cells (Sytox blue$^-$) not shown. (**A**) Sort gates of erythrocyte progenitors (EryP II and EryP III) from bone marrow. (**B**) Sort gates for HSC (LT+ST), MPP from LSK compartment (upper right), and CMP, GMP, MEP from LK compartment (lower right) from bone marrow. (**C**) Sort gates for CLP in bone marrow. (**D**) Sort gates for proB cells (fraction (Fra) B and Fr. C) in bone marrow. (**E**) Sort gates of CD4$^+$ or CD8$^+$ T cells from spleen. (**F**) Sort gates of granulocytes from bone marrow. (**G** and **H**) Sort gates of conventional B (B2) cells (G), and monocytes (H) from spleen. (**I**) Sort gates of preT cells (DN2 and DN3) from the thymus. See Materials and Methods for detailed antibody list and marker phenotypes.

**Fig. S6. Barcode correlation in independent sample repeats in vivo**. (**A**) Scatter plots compare barcode abundance between indicated RNA sample replicates analyzed in single cells in each experiment; Venn diagrams show numbers of unique and shared barcodes between the RNA sample replicates. Barcode abundance levels were quantified by counting single cells that expressed the specific barcodes. (**B**) Scatter plots compare barcode abundance between indicated DNA sample replicates analyzed in bulk in each experiment; Venn diagrams show numbers of unique and shared barcodes between the DNA sample replicates. Barcode abundance levels were quantified using read counts of individual barcodes. (**C**) Scatter plots compare barcode abundance between RNA and DNA samples of indicated cell populations in each experiment; Venn diagrams show numbers of unique and shared barcodes between the RNA and DNA samples. RNA samples were analyzed in single cells and the barcode abundance levels were quantified by counting single cells that expressed the specific barcodes, whereas DNA samples were analyzed in bulk and the barcode abundance levels were quantified using read counts of individual barcodes. For scatter plots in (A), (B), (C), each dot represents a *Polylox* barcode. Sample-specific barcodes are shown in grey background and corresponding percentages were labeled. Blue dashed lines indicate 95% probability bounds of Poisson counting errors. Spearman rank correlation coefficients are indicated at the top of each plot with 95% confidence bounds computed by bootstrap.

**Fig. S7. Barcodes in HSC, progenitor cells and peripheral lineages**. (**A**) Heatmaps showing presence of barcodes in peripheral lineages (rows, rare barcodes with *Pgen* $< 5 \times 10^{-4}$ and detected in two independent sample replicates; columns, cell populations). Barcode counts obtained from sample replicates were added up. Frequencies of barcodes are represented by the color-coded scales on the right. (**B**) Hierarchical clustering of barcode rank correlations between the peripheral lineages shown in (A) for lineage reconstruction. Spearman rank correlation coefficients ($\rho$) are represented by the color-coded scales on the right. The dendrograms obtained by hierarchical clustering indicate a major split between myelo-erythroid and lymphoid lineages. (**C** and **D**) Heatmaps of barcode presence in HSC (first lane), MPP (second lane), progenitors, and downstream erythroid, myeloid and lymphoid lineages as indicated. Barcodes were analysed in single cells or bulk as indicated. Based on the presence of HSC barcodes in peripheral lineages, HSC clones were classified into inactive, myelo-erythroid-restricted, myelo-erythroid biased and multilineage. Color scale indicates barcode frequencies; barcodes in black ink have $P_{gen} < 5 \times 10^{-4}$, barcodes in red ink have $P_{gen} > 5 \times 10^{-4}$. Gr, granulocytes; EryP, erythroid-progenitors stage II/III; Mono, Monocytes; HSC, hematopoietic stem cell; MPP, multipotent progenitor; CMP, common myeloid progenitor; GMP, granulocyte-monocyte progenitor; MEP, megakaryocyte-erythrocyte progenitor; CLP, common lymphoid progenitor.

**Fig. S8. Projection of fate-defined HSC clones onto the transcriptional landscape of hematopoiesis.** (**A**, **E**, and **I**) Transcriptional landscape of phenotypically defined cell populations in Exp. 2 (A), Exp. 3 (E) and Exp. 4 (I) reconstructed using the diffusion map. Each dot represents a single cell of the indicated cell populations. (**B**, **F**, and **J**) Identification of branches in the transcriptional landscapes using diffusion pseudotime. Lineage branches color-coded and labeled according to marker gene expression. (**C**, **G**, and **K**) Pseudo-temporal ordering of cells according to pseudotime rank. (**D**, **H**, and **L**) Projection of fate-defined hematopoietic stem and progenitor cells onto the transcriptional landscape. In individual plots, cells derived from the indicated HSC clone, and colored according to their phenotypically defined stem and progenitor stages, are located on the landscape (grey background). Examples of inactive (green frame), myelo-erythroid-restricted (blue frame), and multilineage clones (red frame), are shown.

**Fig. S9. Heatmap of marker gene expression along diffusion pseudotime rank of cells in the common stem and lineage branches of the transcriptional landscape.** Color scale indicates normalized gene expression values. Data are from Exp. 1. Stem, hematopoietic stem and progenitor trunk; E, erythroid; Mk, megakaryocytic; Gr, granulocytic; Lym/ly, lymphoid; B, B cell.

**Fig. S10. HSC in the same fate category are transcriptionally similar.** Pseudotime rank variability of inactive HSC (n = 21, from Exp. 1 - 3), myelo-erythroid-restricted HSC (n = 67, from Exp. 1 - 3), and active (i.e. multilineage) HSC (n = 100, from Exp. 1 - 3) was measured as interquartile range and shown by colored arrows in each panel. For each fate category, pseudotime rank variability was compared to the null distribution (shown as grey histogram) of interquartile range of pseudotime ranks, obtained by randomly drawing the same number of HSC 10,000 times. The empirical p-values were computed as the number of interquartile range values obtained by random draws that are smaller than or equal to the observed interquartile range value for HSC from the given fate category, normalized by the total number of random draws.

**Fig. S11. Further transcriptome analyses of inactive and active HSC clones**. (**A**) Data of Fig. 3E, but shown as violin plot, showing similar distributions of cell-cycle scores in inactive (n = 24) and active multilineage (n = 103) HSC. (**B**) Heatmap showing the monotonic expression changes of 30 genes (y-axis) exhibiting the most significant dependence on the pseudotime progression (x-axis). All HSC with transcriptomes profiled in Exp. 1 - 3 are shown. (C) As in B, but only the subset of fate-defined HSC is shown (n = 21 inactive and n = 100 active / multilineage HSC from Exp. 1 - 3).

**Fig. S12. Transcriptome-based supervised classification of HSC fates**. (**A**) Receiver operating characteristic (ROC) curve showing the performance of supervised random forest classification (Materials and Methods) of HSC to inactive or active multilineage fates based on transcriptome data. (**B**) ROC curve showing the performance of random forest classification of HSC to myelo-erythroid restricted or multilineage fates based on transcriptome data. (**C**) ROC curve showing the performance of random forest classification of MPP to myelo-erythroid restricted or multilineage fates based on transcriptome data. The dashed diagonal lines indicate performance of random classifiers, corresponding to an AUC = 0.5. AUC, area under curve.

### Fig. S1

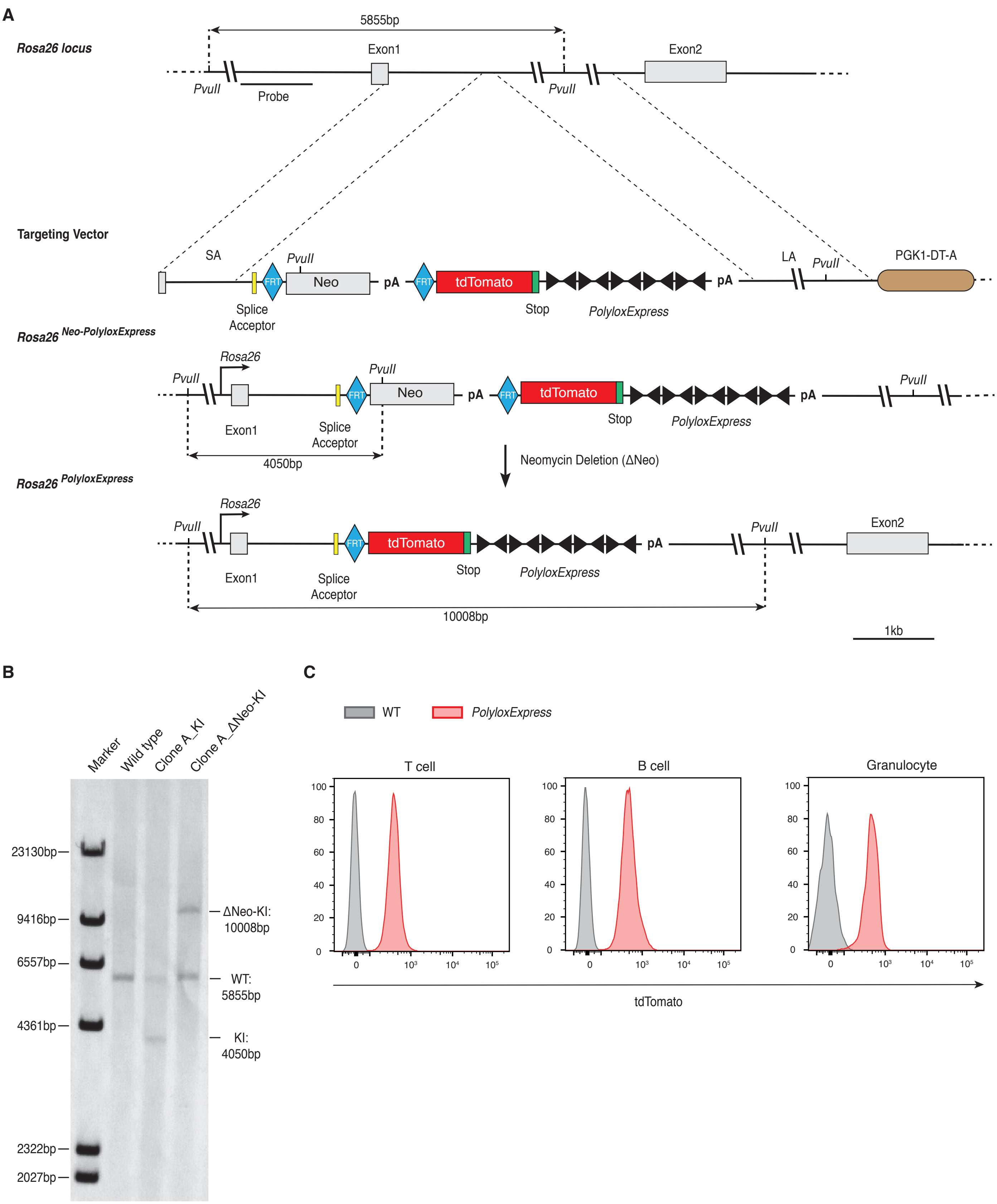
A
Rosa26 locus
5855bp
Exon1
Exon2
PvuII
Probe
PvuII
Targeting Vector
SA
PvuII
Neo
pA
FRT
tdTomato
pA
LA
PvuII
PGK1-DT-A
Splice Acceptor
Stop
PolyloxExpress
Rosa26 Neo-PolyloxExpress
Rosa26
PvuII
PvuII
Neo
pA
tdTomato
pA
PvuII
Exon1
Splice Acceptor
Stop
PolyloxExpress
4050bp
Neomycin Deletion (ΔNeo)
Rosa26 PolyloxExpress
Rosa26
PvuII
tdTomato
pA
PvuII
Exon2
Exon1
Splice Acceptor
Stop
PolyloxExpress
10008bp
1kb
B
Marker
Wild type
Clone A_KI
Clone A_ΔNeo-KI
23130bp
9416bp
6557bp
4361bp
2322bp
2027bp
ΔNeo-KI: 10008bp
WT: 5855bp
KI: 4050bp
C
WT
PolyloxExpress
T cell
B cell
Granulocyte
100
80
60
40
20
0
0
10³
10⁴
10⁵
tdTomato

**Fig. S2**

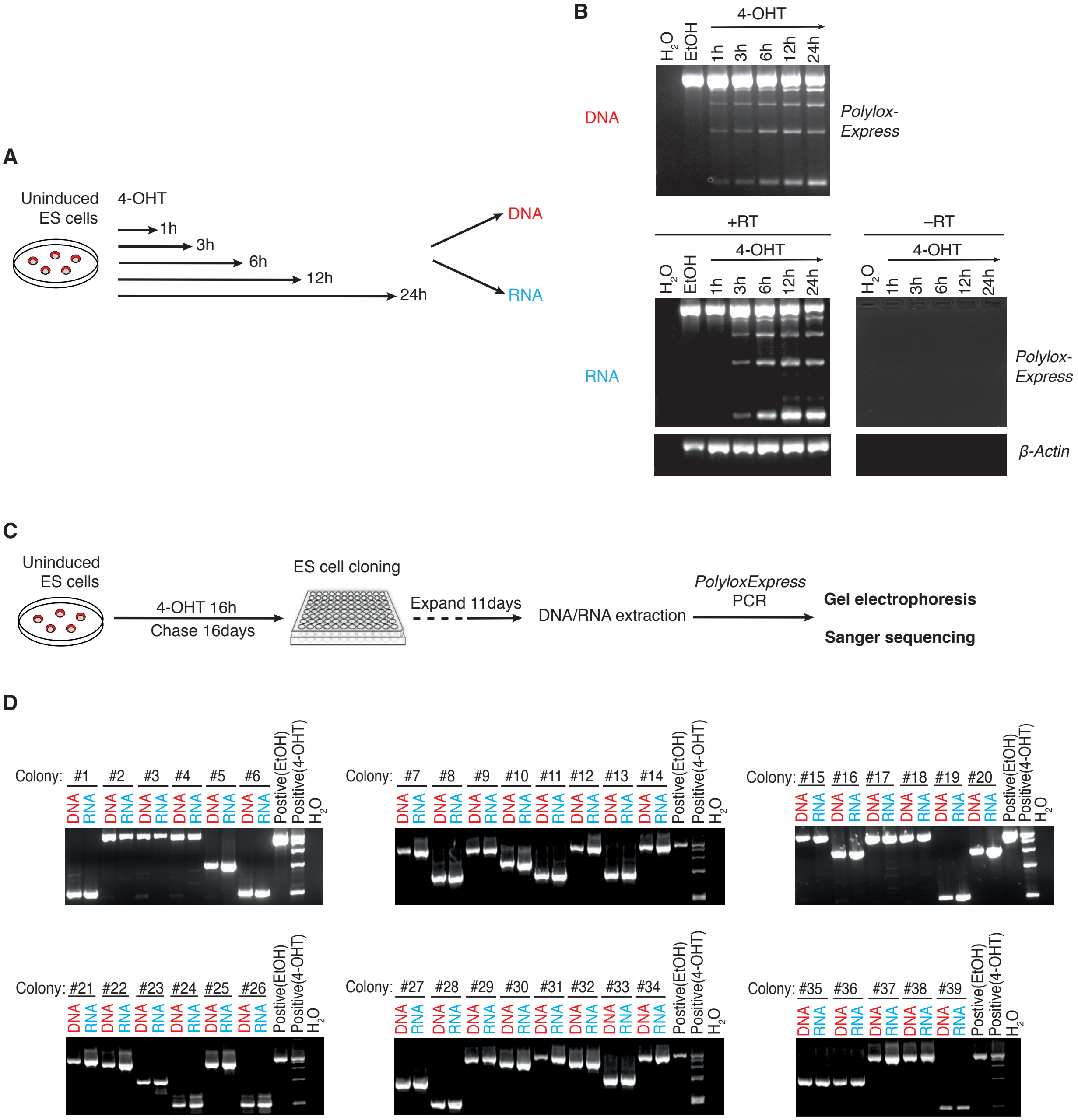
A
Uninduced ES cells
4-OHT
1h
3h
6h
12h
24h
DNA
RNA
B
DNA
RNA
4-OHT
H2O
EtOH
1h
3h
6h
12h
24h
Polylox-Express
+RT
–RT
β-Actin
C
Uninduced ES cells
4-OHT 16h
Chase 16days
ES cell cloning
Expand 11days
DNA/RNA extraction
PolyloxExpress PCR
Gel electrophoresis
Sanger sequencing
D
Colony:
DNA
RNA
Postive(EtOH)
Positive(4-OHT)
H2O

#### Fig. S3

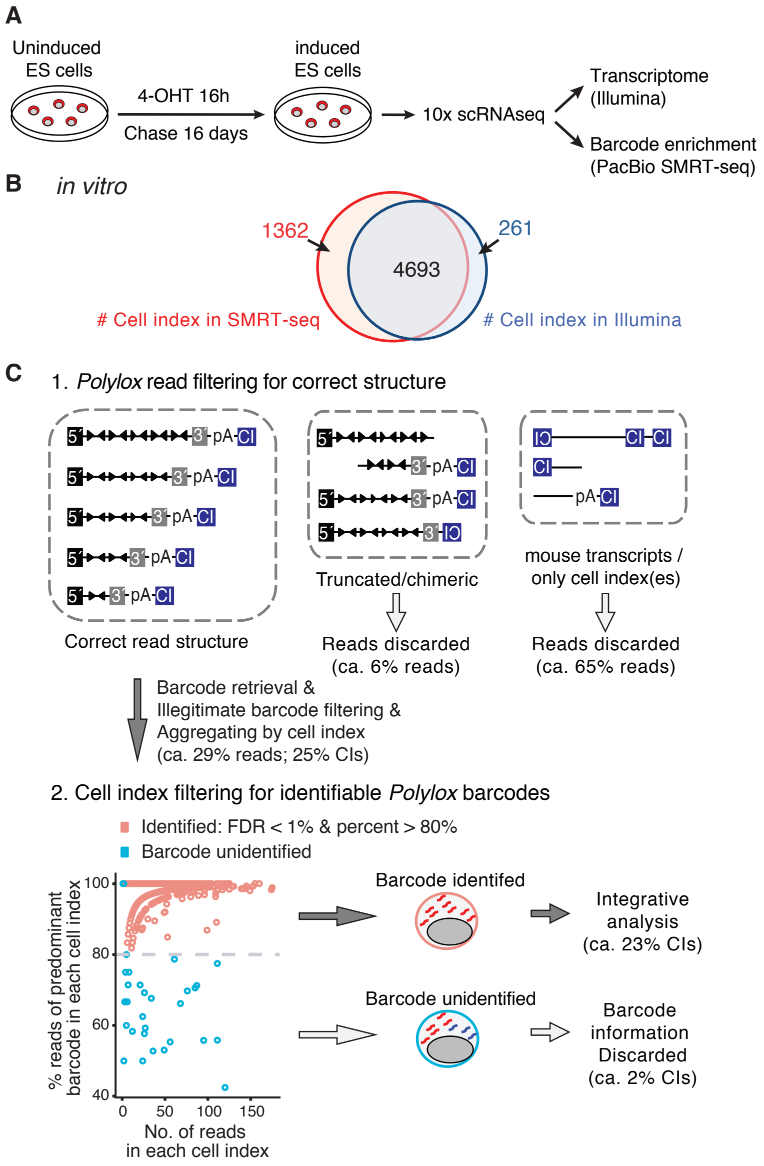
A
Uninduced ES cells
4-OHT 16h
Chase 16 days
induced ES cells
10x scRNAseq
Transcriptome (Illumina)
Barcode enrichment (PacBio SMRT-seq)
B
in vitro
1362
4693
261
### Cell index in SMRT-seq
### Cell index in Illumina
C
1. Polylox read filtering for correct structure
Correct read structure
Truncated/chimeric
Reads discarded (ca. 6% reads)
mouse transcripts / only cell index(es)
Reads discarded (ca. 65% reads)
Barcode retrieval & Illegitimate barcode filtering & Aggregating by cell index (ca. 29% reads; 25% CIs)
2. Cell index filtering for identifiable Polylox barcodes
Identified: FDR < 1% & percent > 80%
Barcode unidentified
% reads of predominant barcode in each cell index
No. of reads in each cell index
Barcode identifed
Integrative analysis (ca. 23% CIs)
Barcode unidentified
Barcode information Discarded (ca. 2% CIs)

#### Fig. S4

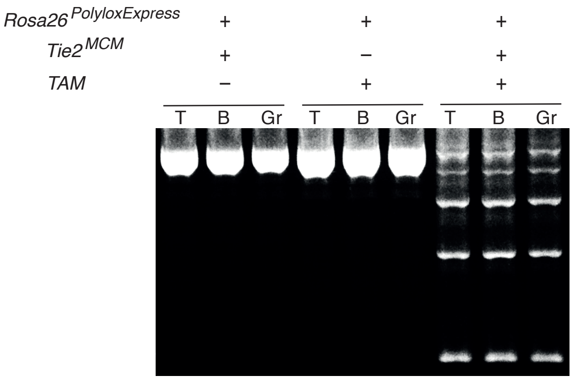

#### Fig. S5

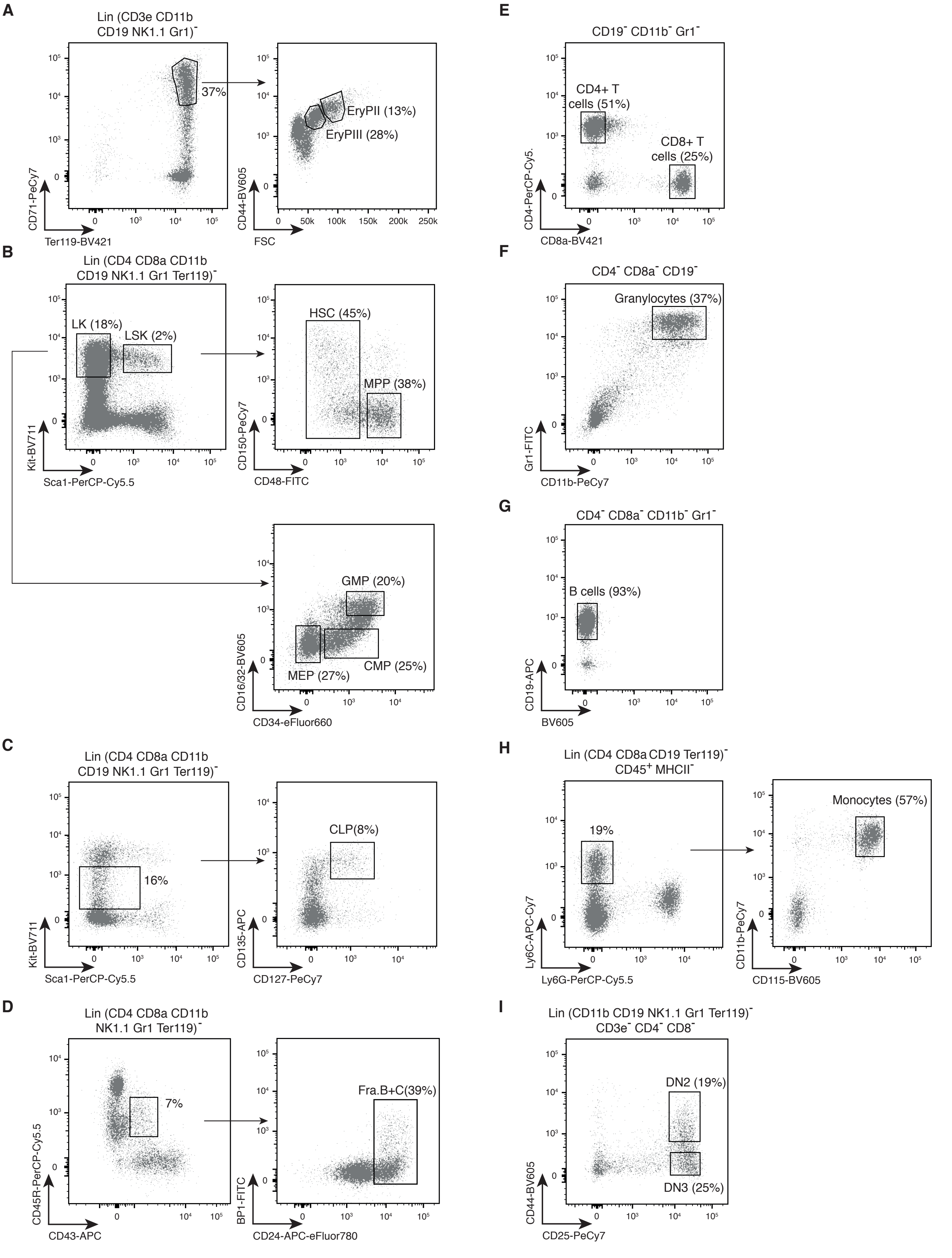

A
Lin (CD3e CD11b CD19 NK1.1 Gr1)⁻
37%
EryPII (13%)
EryPIII (28%)
CD71-PeCy7
Ter119-BV421
CD44-BV605
FSC
B
Lin (CD4 CD8a CD11b CD19 NK1.1 Gr1 Ter119)⁻
LK (18%)
LSK (2%)
HSC (45%)
MPP (38%)
GMP (20%)
CMP (25%)
MEP (27%)
Kit-BV711
Sca1-PerCP-Cy5.5
CD150-PeCy7
CD48-FITC
CD16/32-BV605
CD34-eFluor660
C
Lin (CD4 CD8a CD11b CD19 NK1.1 Gr1 Ter119)⁻
16%
CLP(8%)
Kit-BV711
Sca1-PerCP-Cy5.5
CD135-APC
CD127-PeCy7
D
Lin (CD4 CD8a CD11b NK1.1 Gr1 Ter119)⁻
7%
Fra.B+C(39%)
CD45R-PerCP-Cy5.5
CD43-APC
BP1-FITC
CD24-APC-eFluor780
E
CD19⁻ CD11b⁻ Gr1⁻
CD4+ T cells (51%)
CD8+ T cells (25%)
CD4-PerCP-Cy5.
CD8a-BV421
F
CD4⁻ CD8a⁻ CD19⁻
Granylocytes (37%)
Gr1-FITC
CD11b-PeCy7
G
CD4⁻ CD8a⁻ CD11b⁻ Gr1⁻
B cells (93%)
CD19-APC
BV605
H
Lin (CD4 CD8a CD19 Ter119)⁻ CD45⁺ MHCII⁻
19%
Monocytes (57%)
Ly6C-APC-Cy7
Ly6G-PerCP-Cy5.5
CD11b-PeCy7
CD115-BV605
I
Lin (CD11b CD19 NK1.1 Gr1 Ter119)⁻ CD3e⁻ CD4⁻ CD8⁻
DN2 (19%)
DN3 (25%)
CD44-BV605
CD25-PeCy7

### Fig. S6

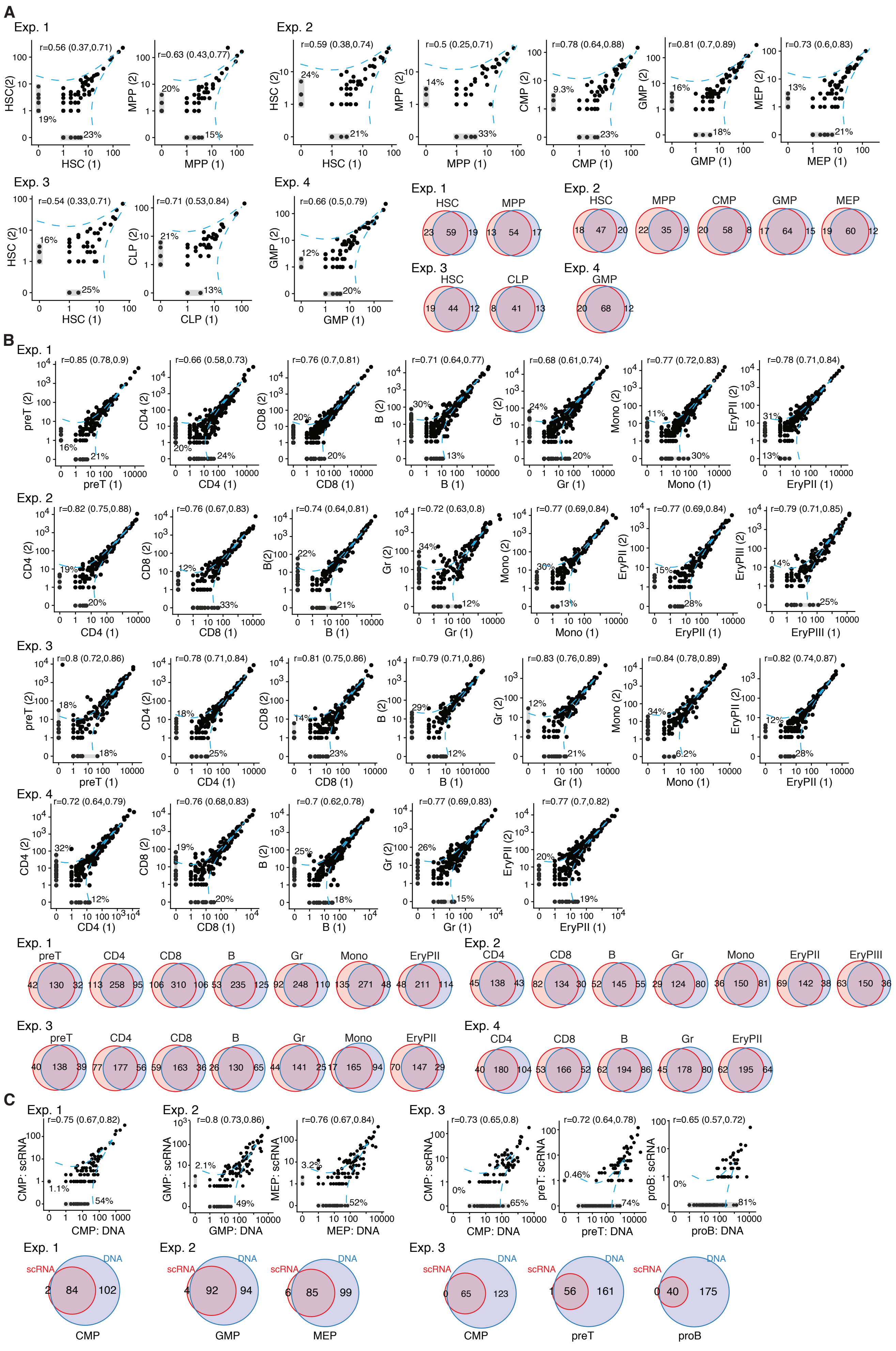

### Fig. S7

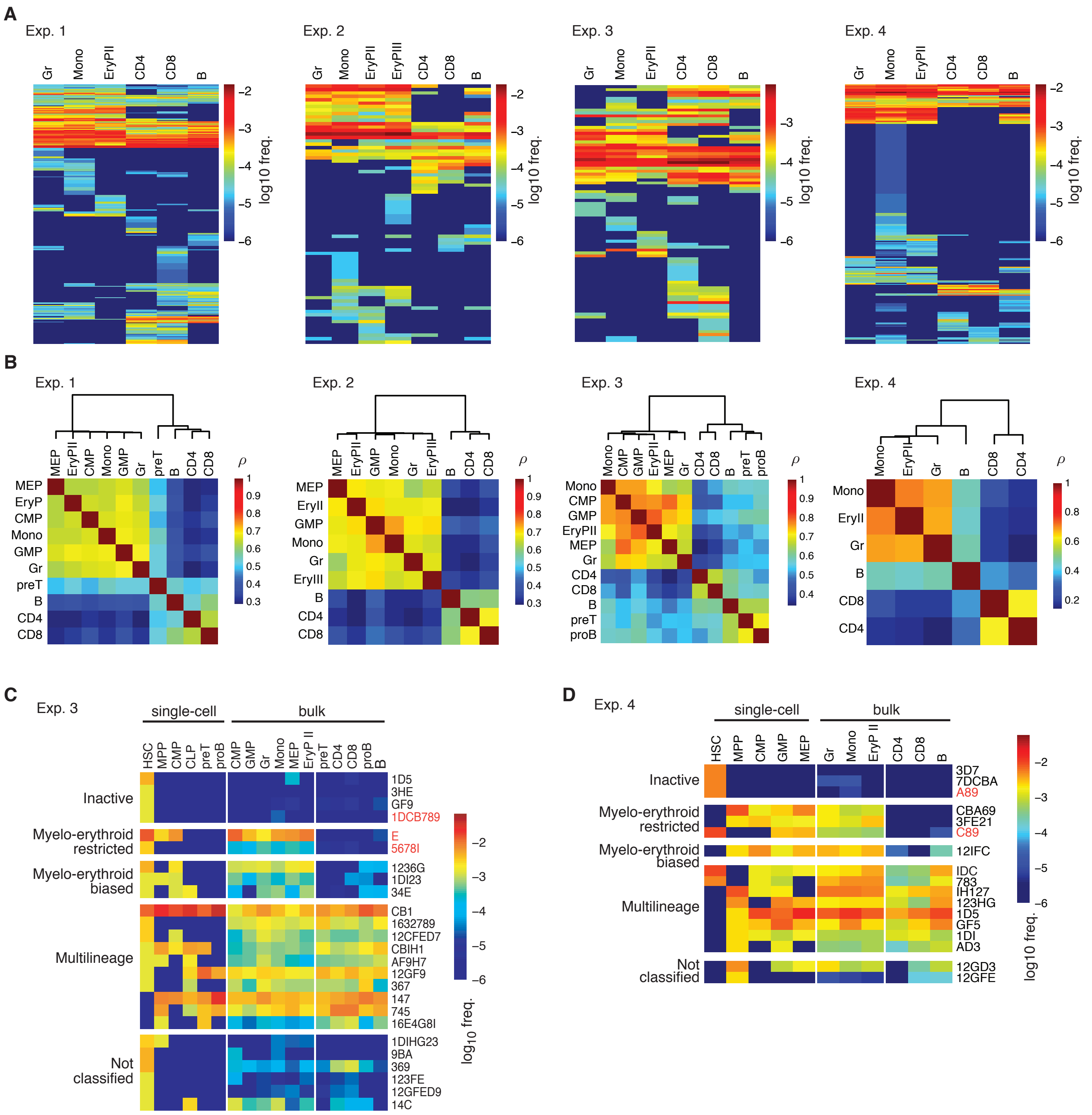
A
Exp. 1
Gr
Mono
EryPII
CD4
CD8
B
log10 freq.
Exp. 2
EryPIII
Exp. 3
Exp. 4
B
ρ
MEP
EryP
CMP
GMP
preT
EryII
EryIII
proB
C
Exp. 3
single-cell
bulk
HSC
MPP
CLP
EryP II
Inactive
Myelo-erythroid restricted
Myelo-erythroid biased
Multilineage
Not classified
1D5
3HE
GF9
1DCB789
E
5678I
1236G
1DI23
34E
CB1
1632789
12CFED7
CBIH1
AF9H7
12GF9
367
147
745
16E4G8I
1DIHG23
9BA
369
123FE
12GFED9
14C
log10 freq.
D
Exp. 4
3D7
7DCBA
A89
CBA69
3FE21
C89
12IFC
IDC
783
IH127
123HG
GF5
1DI
AD3
12GD3
12GFE

Fig. S8

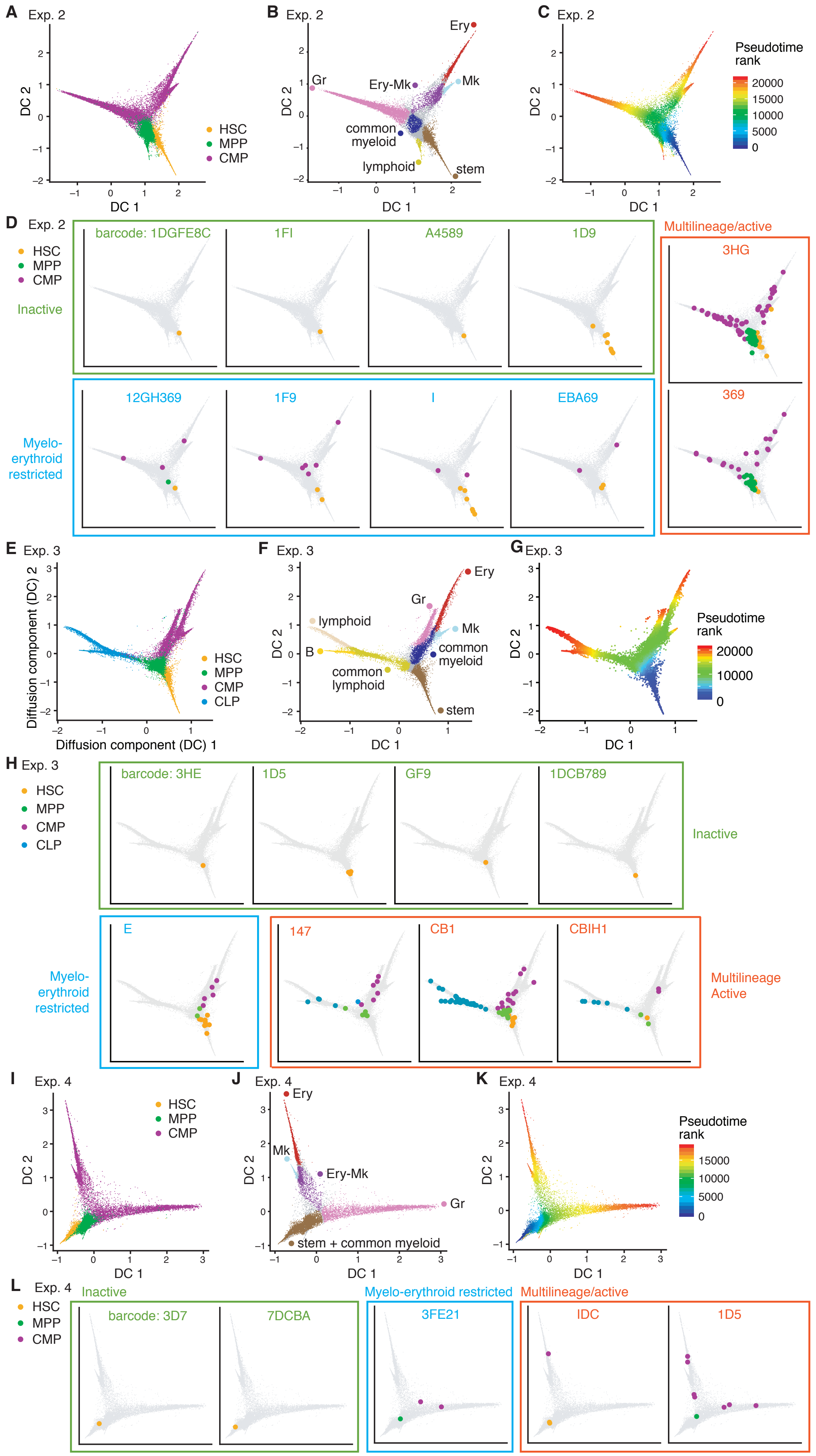

A Exp. 2
HSC
MPP
CMP
DC 1
DC 2
B Exp. 2
Ery
Gr
Ery-Mk
Mk
common
myeloid
lymphoid
stem
C Exp. 2
Pseudotime
rank
20000
15000
10000
5000
0
D Exp. 2
Inactive
barcode: 1DGFE8C
1FI
A4589
1D9
Multilineage/active
3HG
369
Myelo-
erythroid
restricted
12GH369
1F9
I
EBA69
E Exp. 3
Diffusion component (DC) 2
Diffusion component (DC) 1
CLP
F Exp. 3
lymphoid
B
common
lymphoid
G Exp. 3
H Exp. 3
barcode: 3HE
1D5
GF9
1DCB789
E
147
CB1
CBIH1
Multilineage
Active
I Exp. 4
J Exp. 4
stem + common myeloid
K Exp. 4
L Exp. 4
barcode: 3D7
7DCBA
Myelo-erythroid restricted
3FE21
IDC
1D5

**Fig. S9**

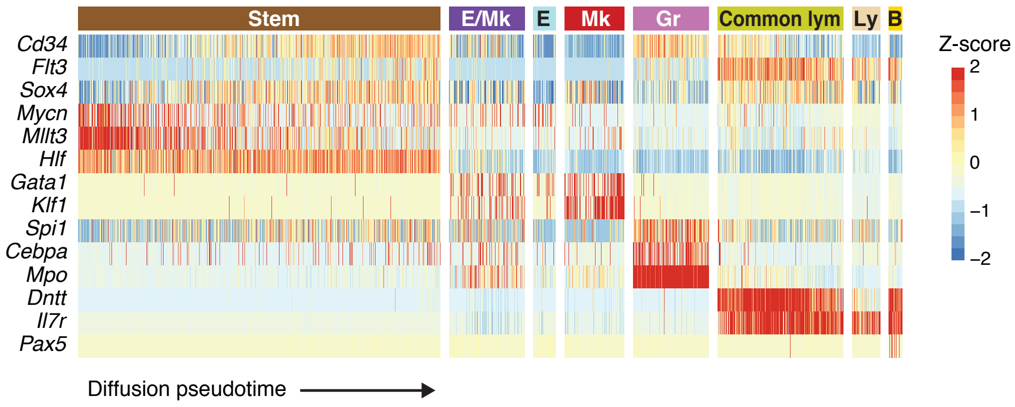
Stem
E/Mk
E
Mk
Gr
Common lym
Ly
B
Cd34
Flt3
Sox4
Mycn
Mllt3
Hlf
Gata1
Klf1
Spi1
Cebpa
Mpo
Dntt
Il7r
Pax5
Z-score
2
1
0
−1
−2
Diffusion pseudotime

#### Fig. S10

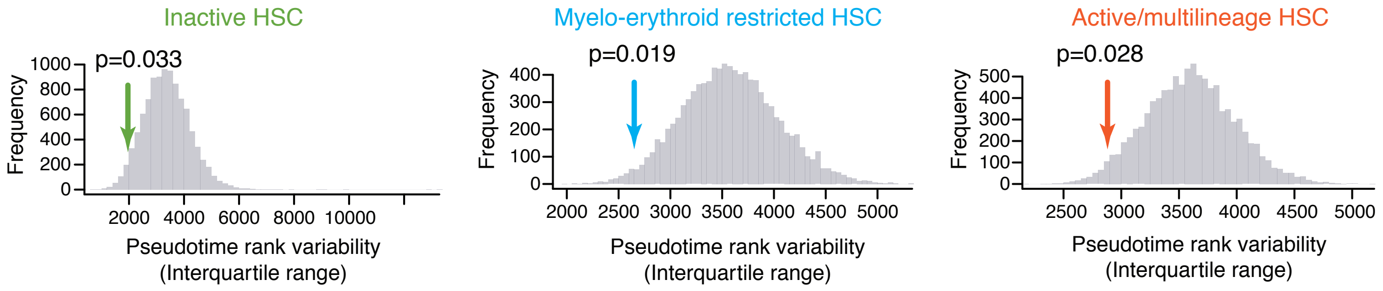

Inactive HSC
p=0.033
Frequency
1000
800
600
400
200
0
2000
4000
6000
8000
10000
Pseudotime rank variability
(Interquartile range)
Myelo-erythroid restricted HSC
p=0.019
Frequency
400
300
200
100
0
2000
2500
3000
3500
4000
4500
5000
Pseudotime rank variability
(Interquartile range)
Active/multilineage HSC
p=0.028
Frequency
500
400
300
200
100
0
2500
3000
3500
4000
4500
5000
Pseudotime rank variability
(Interquartile range)

#### Fig. S11

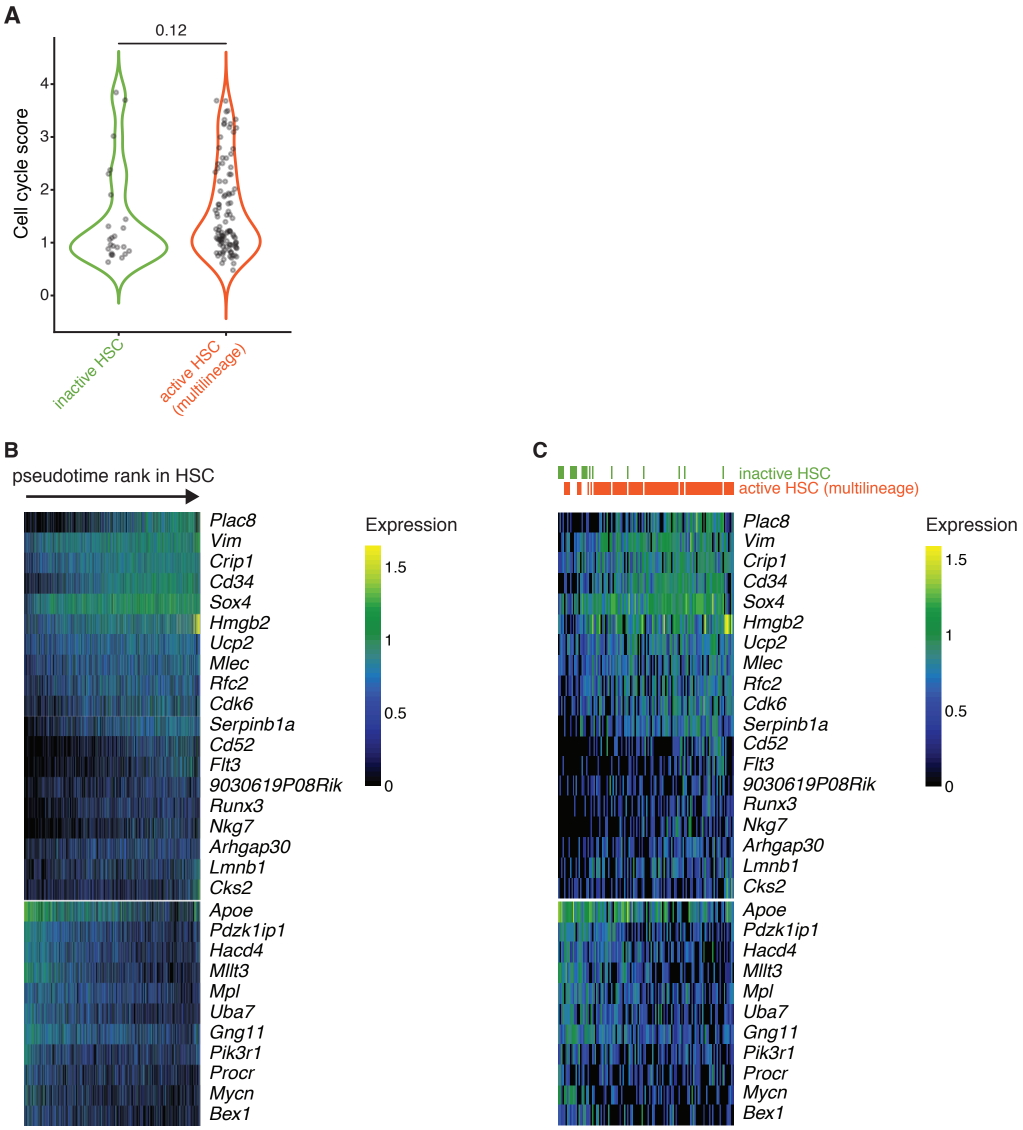

A
0.12
Cell cycle score
4
3
2
1
0
inactive HSC
active HSC (multilineage)
B
pseudotime rank in HSC
C
inactive HSC
active HSC (multilineage)
Plac8
Vim
Crip1
Cd34
Sox4
Hmgb2
Ucp2
Mlec
Rfc2
Cdk6
Serpinb1a
Cd52
Flt3
9030619P08Rik
Runx3
Nkg7
Arhgap30
Lmnb1
Cks2
Apoe
Pdzk1ip1
Hacd4
Mllt3
Mpl
Uba7
Gng11
Pik3r1
Procr
Mycn
Bex1
Expression
1.5
1
0.5
0

**Fig. S12**

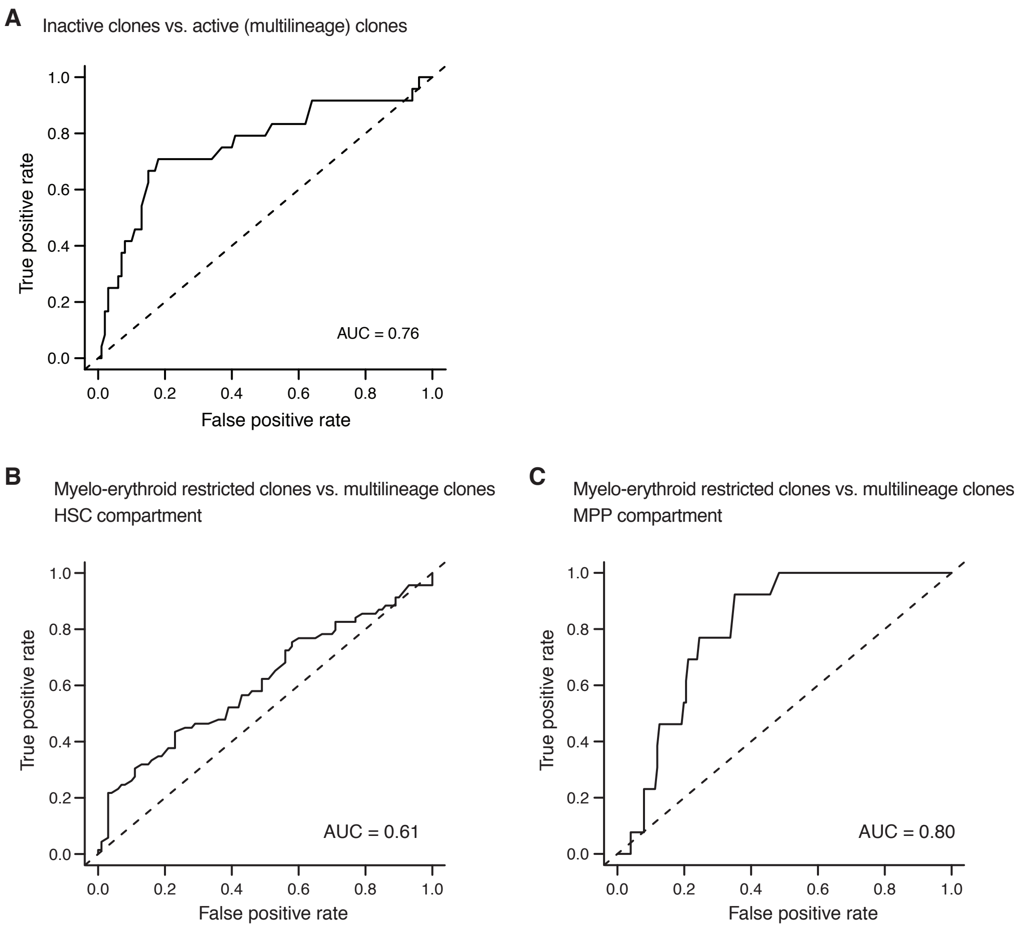
A
Inactive clones vs. active (multilineage) clones
True positive rate
False positive rate
AUC = 0.76
B
Myelo-erythroid restricted clones vs. multilineage clones
HSC compartment
True positive rate
False positive rate
AUC = 0.61
C
Myelo-erythroid restricted clones vs. multilineage clones
MPP compartment
True positive rate
False positive rate
AUC = 0.80

#### Supplementary Tables

**Table S1. The number of barcode types and barcode induction rate in each of the four experiments.**

| | No. barcode types detected | | | | Barcode induction rate |
|---|---|---|---|---|---|
| | Stem + progenitor + peripheral cells | Stem + progenitor cells | | | |
| | Single-cell + bulk | Single-cell + bulk | Single-cell analysis | Bulk analysis | Bulk analysis |
| **Exp. 1** | 1104 | 502 | 133 | 500 | 99.5% |
| **Exp. 2** | 532 | 246 | 133 | 232 | 85.6% |
| **Exp. 3** | 629 | 384 | 111 | 281 | 86.1% |
| **Exp. 4** | 696 | n.a. | 93 | n.a. | 82.1% |
| **Average** | 740 | 377 | 118 | 338 | 88.3% |
| **S.D.** | 252 | 128 | 19 | 143 | 7.6% |

**Table S2. Detailed mouse information in this study.**

| Mouse ID | Exp. ID | Genotype | Sex | Treatment (Age) | Analysis (Age) | Used in figures |
|---|---|---|---|---|---|---|
| TPE#13 | Exp. 1 | *RosaPolyExpressTie2MCM* | F | 1 x TAM (E9.5) | 20 weeks | Fig. 2A-E; Fig. 3B-G; Fig. 4A-J; fig. S5A,C-I; fig. S6A-C; fig. S7A-B; fig. S9; fig. S10; fig. S11; fig. S12 |
| TPE#4 | Exp. 2 | *RosaPolyExpressTie2MCM* | F | 1 x TAM (E9.5) | 17 weeks | Fig. 2A; Fig. 3A-G; Fig. 4A-L; Fig. S5B; fig. S6A-C; fig. S7A-B; fig. S8A-D; fig. S10; fig. S11; fig. S12 |
| TPE#6 | Exp. 3 | *RosaPolyExpressTie2MCM* | M | 1 x TAM (E9.5) | 19 weeks | Fig. 3A-G; Fig. 4A-J; fig. S6A-C; fig. S7A-C; fig. S8E-H; fig. S10; fig. S11; fig. S12 |
| TPE#9 | Exp. 4 | *RosaPolyExpressTie2MCM* | M | 1 x TAM (E9.5) | 7 weeks | Fig. 3B-G; Fig. 4A-I; fig. S6A-B; fig. S7A,B,D; fig. S8I-L; fig. S11A; fig. S12 |
| N.A. | | *Wild type* | N.A. | no treatment | N.A. | fig. S1B |
| #51824 | | *RosaNeo-PolyExpress* | F | no treatment | 7 months | fig. S1B |
| #51823 | | *RosaPolyExpress* | F | no treatment | 7 months | fig. S1B |
| #41068 | | *Wild type* | F | no treatment | 19 weeks | fig. S1C |
| #54754 | | *RosaPolyExpressTie2MCM* | F | no treatment | 15 weeks | fig. S1C |
| #54754 | | *RosaPolyExpressTie2MCM* | F | no treatment | 15 weeks | fig. S4 |
| #50 | | *RosaPolyExpress* | M | 1 x TAM (E9.5) | 18 weeks | fig. S4 |
| #52 | | *RosaPolyExpressTie2MCM* | M | 1 x TAM (E9.5) | 18 weeks | fig. S4 |

**Table S3. Lists of mark genes used to annotate diffusion map branches.**

| Trunk/branch | Marker genes |
|---|---|
| **Stem** | Hlf, Ifitm1, Ly6a |
| **Mk** | Pf4, Vwf, Itga2b |
| **Ery** | Mki67, Gata1, Car2, Hbb-bs, Hba-a2, Car1, Apoe, Gata2 |
| **Mono** | F13a1, Ly86, Csf1r, C1qb |
| **Gr** | Ifitm1, Mpo, Elane, Cebpe, Prss34, Prg2, Gstm1, Fcnb, Ltf, Gfi1, Mmp8, Itgam, Il1b, Ccl6 |
| **Lym** | Dntt, Satb1, Flt3, Ly6d, Rag1, Ebf1, Cd79a, Gata3, Pax5, Vpreb1, Vpreb2, Vpreb3, Igll1, Fcrla, Ccl5, Ncr1, Cd3d |
| **B** | Pax5, Vpreb1, Vpreb2, Vpreb3, Igll1, Fcrla |

**Table S4. Signature gene lists used in this study.**

| S-phase* | G2/M-phase* | LT-HSC | ST-HSC | Dormant HSC[#] | CMP | CLP |
|---|---|---|---|---|---|---|
| Ung | Nuf2 | Apoe | H2afy | Rab44 | Adgrg1 | 2810417H13Rik |
| Usp1 | Cks1b | Rpl9-ps6 | Cd34 | Igf2bp2 | Aldoa | Arpc5l |
| Dscc1 | Cdc20 | Gm10036 | Tmsb10 | Ahnak | Angpt1 | Arpp21 |
| Cdca7 | Gas2l3 | Pdzk1ip1 | Plac8 | Mpo | Ap3s1 | Atp1b1 |
| Mcm6 | Ect2 | Clec1a | Flt3 | Plac8 | Apoe | Atp1b3 |
| Pold3 | Ctcf | Bex4 | Ramp1 | Fam101b | Asap1 | Atp2b4 |
| Pcna | Nusap1 | Sult1a1 | Cd52 | Ablim1 | Atpif1 | B3gnt2 |
| Ubr7 | Kif20b | Aldh1a1 | Serpinb1a | Milr1 | Bin2 | BC035044 |
| Pola1 | Cdca2 | Cavin2 | Sh3bgrl3 | Meg3 | Calr | Bex6 |
| Atad2 | Ckap2 | Mllt3 | Sox4 | Ms4a6c | Car1 | Bfsp2 |
| Wdr76 | Tubb4b | Gm11808 | Gm5111 | Cd53 | Car2 | Blk |
| Dtl | Lbr | Rps21 | mt-Nd4l | Cd7 | Ccl9 | Blnk |
| Hells | Cenpa | Tgm2 | Emb | Hp | Cd34 | Bst2 |
| Gins2 | Kif2c | Nupr1 | Wfdc17 | Tyrobp | Cd63 | Camk1d |
| Fen1 | Hmgb2 | Rpl21 | Uba52 | Raph1 | Cd9 | Ccnd3 |
| Rrm2 | Tmpo | Rps28 | Cd53 | Depdc1a | Clec4d | Cd37 |
| Uhrf1 | Kif23 | Nt5c3 | Gmfg | Glcci1 | Cpa3 | Cd52 |
| E2f8 | Cbx5 | Slamf1 | Cd37 | F13a1 | Creg1 | Cd53 |
| Rpa2 | Ckap5 | Selenom | Crip1 | A930006K02Rik | Csrp3 | Cd72 |
| Mcm5 | Nek2 | Slfn2 | Ctla2a | A330023F24Rik | Ctla2a | Cd79a |
| Blm | Cdk1 | Meg3 | Tespa1 | Sorl1 | Ctsg | Cd93 |
| Brip1 | Smc4 | Rpl38 | Adgrl4 | Irf8 | Dach1 | Cdkn1a |
| Tipin | Tpx2 | Cracr2b | Cmtm7 | Xdh | Dapp1 | Cdv3 |
| Rad51ap1 | Dlgap5 | Uba7 | Gcnt2 | Anxa2 | Elane | Chchd10 |
| Msh2 | Anln | Rps12-ps3 | Hlf | Ccr2 | Etfb | Clec12a |
| Rfc2 | Mki67 | Trim47 | Gpx1 | Card9 | F630028O10Rik | Clec2d |
| Rrm1 | G2e3 | Mmrn1 | Nkg7 | Flrt1 | Fam46a | Cmah |
| Clspn | Tacc3 | Cd9 | Irf2bp2 | Hdc | Fermt3 | Cnn3 |
| Exo1 | Cenpf | Gda | Cd27 | Mmp9 | Gata2 | Cnp |
| Cdc6 | Birc5 | Obscn | Ddx4 | Camp | Gclm | Coro1a |
| Rad51 | Top2a | Ryk | Rabgap1l | Hbb-bt | Gse1 | Cox6a2 |
| Casp8ap2 | Cdca8 | Gm2000 | Taldo1 | Nucb2 | Gstm5 | Crip1 |
| Nasp | Ncapd2 | Rps27 | Shisa5 | Elane | Hdgf | Ctr9 |
| Mcm2 | Anp32e | Kazald1 | Slc16a11 | Amica1 | Hlf | Ctsb |
| Gmnn | Hmmr | Gng11 | BC035044 | Kcnh7 | Hmgb3 | Ctss |
| Ccne2 | Hjurp | Socs2 | Ccl3 | Ifitm6 | Ifitm1 | Dntt |
| Cdc45 | Ckap2l | Rpl7a-ps5 | Lat2 | Cfp | Ifitm2 | Drc7 |
| Chaf1b | Aurkb | Gm10073 | Itgb5 | Hvcn1 | Ifitm3 | Ebf1 |
| Prim1 | Cks2 | Bex1 | Ighm | Smarca5-ps | Itga2b | Egfl7 |
| Tyms | Bub1 | Hoxb2 | Ikzf2 | Ccl5 | Khk | Egln1 |
| Slbp | Psrc1 | Cpne8 | Satb1 | Cd52 | Kit | Evl |
| Mcm4 | Ube2c | Art4 | Chd3 | Siglech | Ldha | Flt3 |
| Pcna-ps2 | Aurka | Gata2 | Shisa8 | Lgals3 | Lmo2 | Foxp1 |
| | Cenpe | Ldha | Gm10076 | Rassf4 | Mpo | Gimap6 |
| | Ccnb2 | Cdkn1c | Arhgap30 | Ccr1 | Ms4a3 | Gm1966 |
| | Cdca3 | Fam110c | Vldlr | Ncf1 | Muc13 | Gm4759 |
| | Ttk | Pbx1 | Fam69b | Igf1 | Nfe2 | Gm5111 |
| | Rangap1 | Grb10 | Sec61b | Cd2 | Ninj1 | Gphn |
| | Ndc80 | Gm11361 | Arhgdib | Lrp4 | Nkg7 | Gpr171 |
| | Cdc25c | Rpl13a | Lcp1 | Dbf4 | Nt5c3 | Gpr25 |
| | Kif11 | Nadk2 | Ifi203 | Klrd1 | Pdcd4 | H2-Ob |
| | Gtse1 | Ifitm7 | Gm2a | Lrrc18 | Pf4 | H2afy |
| | | Trpc6 | Hnrnph1 | Ap1s2 | Pgam1 | Hes6 |
| | | Rpl10-ps3 | Cd47 | Fndc3b | Prdx6 | Hist1h2ae |
| | | Nrgn | Dock10 | Ppfia4 | Prkar2b | Ifi203 |
| | | Esam | Tm6sf1 | E2f2 | Prtn3 | Ifi27l2a |
| | | Serpinb6a | Mlec | Col1a1 | Rab38 | Ikzf1 |

| S-phase* | G2/M-phase* | LT-HSC | ST-HSC | Dormant HSC[#] | CMP | CLP |
|---|---|---|---|---|---|---|
| | | Serpina3g | Dhrs3 | Ctss | S100a1 | Il12a |
| | | Ndn | Ctss | Add2 | Sdsl | Il18rap |
| | | Tcf15 | Ptpre | Tifab | Slc22a3 | Il7r |
| | | Gfi1b | Wfdc18 | Rnf39 | Sord | Irf8 |
| | | Eef2 | Zyx | Cd79a | Srgn | Lbh |
| | | Rpl27-ps3 | Map4k4 | Cybb | Tacstd2 | Lck |
| | | Ifitm3 | Plppr3 | Mcemp1 | Unc119 | Lcp1 |
| | | Plxnc1 | Tyrobp | Pld4 | Vamp5 | Lsm3 |
| | | Rps27rt | Spi1 | Sparc | Vamp8 | Lsp1 |
| | | Slc18a2 | Runx2 | Atp1b1 | Ybx3 | Ly6d |
| | | Ppic | Fchsd2 | Wfdc17 | Zfpm1 | Ly6e |
| | | Rhof | Gpr171 | Hba-a2 | | Ly86 |
| | | Cavin3 | Pcgf5 | Vpreb3 | | Maml3 |
| | | Cenpt | Coro1a | Lonrf2 | | Marcks |
| | | Tinagl1 | Dock8 | Pid1 | | Med13l |
| | | Csgalnact1 | Ltb | Ms4a3 | | Mef2c |
| | | Plxdc2 | Laptm5 | Hbb-bs | | Mgst2 |
| | | Rftn1 | Kcnk12 | Lmna | | Mndal |
| | | Clec2d | Sept1 | Kcnd3 | | Mpeg1 |
| | | Ccnd2 | Rgs2 | Cd209a | | Mta3 |
| | | Mycn | Fkbp3 | Yap1 | | Myl10 |
| | | Hacd4 | Runx3 | Ccl6 | | Myl4 |
| | | Txnip | Mgst1 | Itga9 | | Mzb1 |
| | | Tmem40 | Coro7 | Csf2rb2 | | Notch1 |
| | | Cited2 | Anxa2 | Fam196a | | Nucb2 |
| | | Il11ra1 | BC028528 | Lrrc23 | | Parp1 |
| | | Klf9 | Ucp2 | Ect2 | | Pgls |
| | | Khk | Plek | F630028O10Rik | | Pkig |
| | | Ftl1-ps1 | St8sia4 | Dse | | Psap |
| | | Rpl13a-ps1 | Rnase6 | Nova1 | | Pten |
| | | Sord | Btg2 | Pik3r6 | | Ptp4a3 |
| | | Tbxas1 | Ikzf1 | B3galt2 | | Ptprcap |
| | | Rpl36a-ps1 | H2-DMa | Bmp8a | | Rac2 |
| | | Ubl7 | Arhgap15 | Irak2 | | Rag1 |
| | | Cd63 | Ifi27l2a | Klhl35 | | Ramp1 |
| | | Epb41l4b | Cdk6 | Wfdc21 | | Rpgrip1 |
| | | Pdlim1 | Cd44 | Sell | | Runx2 |
| | | Nedd4 | Samsn1 | Tulp1 | | Satb1 |
| | | Wdr89 | Phf14 | Klra13-ps | | Sdc1 |
| | | Sdsl | Top1 | Camk2n1 | | Sdc4 |
| | | Tmem258 | Cd72 | Adam8 | | Sell |
| | | Cd81 | Scd2 | Fcrla | | Selplg |
| | | Adgrg1 | Pan3 | AI607873 | | Sept1 |
| | | Tubb4b | Iqgap1 | Cebpe | | Siglech |
| | | Hsd17b10 | Vim | Zfp82 | | Smad7 |
| | | Slc48a1 | Il17ra | Sema3e | | Smim14 |
| | | Wfdc2 | Osbpl8 | Arc | | Stk17b |
| | | Ifitm1 | Vav3 | Slc4a1 | | Tbxa2r |
| | | Bola2 | Cux1 | Ank1 | | Tcf4 |
| | | Ccnd3 | Slco3a1 | Ctsg | | Tfrc |
| | | Tsc22d3 | Tia1 | Fabp4 | | Tifa |
| | | Mecom | Mgat1 | Mb21d2 | | Tmem108 |
| | | Rbp1 | Fam117a | Upb1 | | Tmem121 |
| | | S100a6 | Il12a | BC035044 | | Tmem173 |
| | | Hmgn3 | Pcp4l1 | Sh2d2a | | Tmem229b |
| | | Fbxo9 | Man1a | Evi2a | | Tmsb10 |
| | | Aplp2 | Egfl7 | Plvap | | Tpm4 |
| | | Nfkbia | Rasgrp2 | Ms4a4b | | Tsc22d1 |
| | | Pnp | Ifi27 | Sfrp2 | | Tspan13 |

| S-phase* | G2/M-phase* | LT-HSC | ST-HSC | Dormant HSC# | CMP | CLP |
|---|---|---|---|---|---|---|
| | | Krt18 | Hist1h2ap | Atp8b4 | | Tspan2 |
| | | Fut8 | Hoxa9 | | | Ttc13 |
| | | Gm19590 | Fos | | | Tyrobp |
| | | Iigp1 | | | | Unc93b1 |
| | | Pik3r1 | | | | Vpreb3 |
| | | Tuba1b | | | | Wasf2 |
| | | AY036118 | | | | Xpc |
| | | | | | | Xrcc6 |

**Table S5. Fate and number of cells of each HSC clone in the four experiments.**

| Exp. 1 | Barcode | # Cells | % Cells | Fate |
|---|---|---|---|---|
| | **total HSC** | 1876 | | |
| HSC clones (barcode Pgen < 5e-4) | 1HED9 | 16 | 0.85% | mye-ery biased |
| | 16G | 15 | 0.80% | multilineage |
| | C49 | 13 | 0.69% | multilineage |
| | 125HI | 5 | 0.27% | multilineage |
| | 1F7 | 5 | 0.27% | mye-ery biased |
| | 3FE | 5 | 0.27% | mye-ery biased |
| | C69 | 5 | 0.27% | multilineage |
| | C6E | 5 | 0.27% | mye-ery restricted |
| | 3HG | 4 | 0.21% | multilineage |
| | ED189 | 4 | 0.21% | multilineage |
| | 1256G | 4 | 0.21% | multilineage |
| | 123FE | 3 | 0.16% | mye-ery biased |
| | 1DE87 | 3 | 0.16% | inactive |
| | 3HED9 | 3 | 0.16% | mye-ery restricted |
| | A45 | 3 | 0.16% | mye-ery restricted |
| | 34A | 2 | 0.11% | multilineage |
| | 369 | 2 | 0.11% | multilineage |
| | 36G | 2 | 0.11% | multilineage |
| | A4GFE89 | 2 | 0.11% | not classified |
| | ABC | 2 | 0.11% | multilineage |
| | 5DCB9 | 2 | 0.11% | multilineage |
| | CB56IHG | 2 | 0.11% | mye-ery biased |
| | GF1 | 2 | 0.11% | mye-ery biased |
| | 1456I | 1 | 0.05% | mye-ery biased |
| | 163B9 | 1 | 0.05% | mye-ery biased |
| | 3F145HI | 1 | 0.05% | mye-ery restricted |
| | CBAHG | 1 | 0.05% | multilineage |
| | GF1HED9 | 1 | 0.05% | mye-ery restricted |
| | IH1 | 1 | 0.05% | inactive |
| | 123FE4G | 1 | 0.05% | mye-ery restricted |
| | 1HE | 1 | 0.05% | not classified |
| | A69 | 1 | 0.05% | multilineage |

| Exp. 2 | Barcode | # Cells | % Cells | Fate |
|---|---|---|---|---|
| | **total HSC** | 942 | | |
| HSC clones (barcode Pgen < 5e-4) | 3HG | 24 | 1.28% | multilineage |
| | 3F9 | 13 | 0.69% | mye-ery biased |
| | 1DCBGFE | 6 | 0.32% | mye-ery restricted |
| | 369 | 6 | 0.32% | multilineage |
| | 14C89 | 5 | 0.27% | mye-ery biased |
| | 1FEDC | 5 | 0.27% | mye-ery restricted |
| | EBA69 | 3 | 0.16% | mye-ery restricted |
| | 1DGFE8C | 2 | 0.11% | inactive |
| | 9812GFEDC | 2 | 0.11% | not classified |
| | 1FCBG8I | 2 | 0.11% | mye-ery biased |
| | 12GFE | 1 | 0.05% | mye-ery restricted |
| | 12GFE49 | 1 | 0.05% | mye-ery restricted |
| | 12GH369 | 1 | 0.05% | mye-ery restricted |
| | 1FI | 1 | 0.05% | inactive |
| | A69 | 1 | 0.05% | inactive |
| | 1234GHI | 1 | 0.05% | multilineage |

| | Barcode | # Cells | % Cells | Fate |
|---|---|---|---|---|
| | 1278EDC | 1 | 0.05% | mye-ery restricted |
| | A4589 | 1 | 0.05% | inactive |

| Exp. 3 | Barcode | # Cells | % Cells | Fate |
|---|---|---|---|---|
| | **total HSC total HSC** | 686 | | |
| HSC clones (barcode Pgen < 5e-4) | CB1 | 8 | 0.43% | multilineage |
| | 1236G | 3 | 0.16% | mye-ery biased |
| | 1D5 | 3 | 0.16% | inactive |
| | 9BA | 3 | 0.16% | not classified |
| | 369 | 3 | 0.16% | not classified |
| | 12CFED7 | 2 | 0.11% | multilineage |
| | 1632789 | 2 | 0.11% | multilineage |
| | 1DIHG23 | 2 | 0.11% | not classified |
| | 12GFED9 | 1 | 0.05% | not classified |
| | 1DI23 | 1 | 0.05% | mye-ery biased |
| | 34E | 1 | 0.05% | mye-ery biased |
| | 3HE | 1 | 0.05% | inactive |
| | AF9H7 | 1 | 0.05% | multilineage |
| | CBIH1 | 1 | 0.05% | multilineage |
| | GF9 | 1 | 0.05% | inactive |
| | 123FE | 1 | 0.05% | not classified |
| | 12GF9 | 1 | 0.05% | multilineage |
| | 14C | 1 | 0.05% | not classified |
| | 367 | 1 | 0.05% | multilineage |

| Exp. 4 | Barcode | # Cells | % Cells | Fate |
|---|---|---|---|---|
| | **total HSC** | 201 | | |
| HSC clones (barcode Pgen < 5e-4) | IDC | 2 | 0.11% | multilineage |
| | 3D7 | 1 | 0.05% | inactive |
| | 783 | 1 | 0.05% | multilineage |
| | 7DCBA | 1 | 0.05% | inactive |

| clone size | mean | s.d. |
|---|---|---|
| **inactive** | 1.5 | 0.8 |
| **mye-ery restricted** | 2.5 | 1.9 |
| **multilineage** | 4.3 | 5.4 |

**Table S6. Differentially expressed genes.**

DEGs of inactive vs. active (multilineage) HSC

| Gene Symbol | log(fold-change) | P value | adj. P value | Up-regulated in active HSC? |
|---|---|---|---|---|
| Cd34 | -1.07 | 6.75E-06 | 2.79E-03 | Yes |
| Trim47 | 1.07 | 7.47E-06 | 2.79E-03 | No |
| Vim | -1.00 | 2.84E-05 | 7.07E-03 | Yes |
| Mycn | 0.97 | 4.31E-05 | 8.06E-03 | No |
| Uba7 | 0.95 | 6.06E-05 | 9.07E-03 | No |
| Serpinb1a | -0.94 | 7.49E-05 | 9.34E-03 | Yes |
| Cdkn1c | 0.90 | 1.59E-04 | 1.70E-02 | No |
| Gmfg | -0.88 | 2.00E-04 | 1.87E-02 | Yes |
| Sult1a1 | 0.87 | 2.40E-04 | 1.99E-02 | No |
| Hoxb2 | 0.86 | 2.95E-04 | 2.21E-02 | No |
| Tgm2 | 0.84 | 4.16E-04 | 2.83E-02 | No |
| Ccnd2 | 0.82 | 5.54E-04 | 3.46E-02 | No |
| Slfn2 | 0.82 | 6.07E-04 | 3.49E-02 | No |
| Cyp51 | 0.80 | 8.13E-04 | 4.20E-02 | No |
| Gm5111 | -0.79 | 9.25E-04 | 4.20E-02 | Yes |
| Nupr1 | 0.78 | 9.84E-04 | 4.20E-02 | No |
| Stmn1 | -0.78 | 1.05E-03 | 4.20E-02 | Yes |
| Nrgn | 0.78 | 1.05E-03 | 4.20E-02 | No |
| Clec1a | 0.78 | 1.07E-03 | 4.20E-02 | No |
| Bex1 | 0.77 | 1.16E-03 | 4.33E-02 | No |
| Gng11 | 0.76 | 1.30E-03 | 4.63E-02 | No |
| Zfp873 | 0.75 | 1.50E-03 | 5.11E-02 | No |
| Cpne8 | 0.75 | 1.65E-03 | 5.37E-02 | No |
| Pdzk1ip1 | 0.73 | 2.10E-03 | 6.55E-02 | No |
| Ctla2a | -0.72 | 2.40E-03 | 7.18E-02 | Yes |
| Ptma | -0.71 | 2.72E-03 | 7.79E-02 | Yes |
| Adgrl4 | -0.71 | 2.81E-03 | 7.79E-02 | Yes |
| S100a6 | 0.70 | 3.39E-03 | 9.05E-02 | No |
| Esam | 0.69 | 3.68E-03 | 9.49E-02 | No |
| Plac8 | -0.68 | 3.98E-03 | 9.64E-02 | Yes |
| Mmrn1 | 0.68 | 4.03E-03 | 9.64E-02 | No |
| Aplp2 | 0.68 | 4.12E-03 | 9.64E-02 | No |
| Ifitm1 | 0.68 | 4.50E-03 | 1.02E-01 | No |
| Cd53 | -0.67 | 5.06E-03 | 1.11E-01 | Yes |
| Gcnt2 | -0.66 | 5.64E-03 | 1.21E-01 | Yes |
| Bex4 | 0.65 | 6.06E-03 | 1.26E-01 | No |
| Apoe | 0.64 | 6.80E-03 | 1.33E-01 | No |
| Slc25a33 | 0.64 | 6.91E-03 | 1.33E-01 | No |
| Ifi27 | -0.64 | 6.93E-03 | 1.33E-01 | Yes |
| Gas2l3 | 0.63 | 8.35E-03 | 1.56E-01 | No |
| Mt1 | 0.62 | 8.78E-03 | 1.58E-01 | No |
| Fut7 | -0.62 | 8.86E-03 | 1.58E-01 | Yes |
| Grb10 | 0.61 | 9.79E-03 | 1.67E-01 | No |
| Gbp7 | 0.61 | 9.83E-03 | 1.67E-01 | No |
| Erp27 | 0.61 | 1.03E-02 | 1.71E-01 | No |
| Lsp1 | -0.60 | 1.10E-02 | 1.76E-01 | Yes |
| Serpina3g | 0.60 | 1.11E-02 | 1.76E-01 | No |
| Sgms1 | 0.60 | 1.14E-02 | 1.77E-01 | No |
| Fam46a | -0.60 | 1.18E-02 | 1.77E-01 | Yes |
| Med12l | 0.60 | 1.19E-02 | 1.77E-01 | No |
| H2-Eb1 | 0.60 | 1.21E-02 | 1.78E-01 | No |

**DEGs of myelo-erythroid restricted vs. multilineage HSC**

| Gene Symbol | log(fold-change) | P value | adj. P value | Up-regulated in multilineage HSC? |
|---|---|---|---|---|
| Cd34 | -0.75 | 2.24E-06 | 1.48E-03 | Yes |
| H2afy | -0.68 | 1.98E-05 | 6.54E-03 | Yes |
| Serpinb1a | -0.65 | 3.82E-05 | 8.40E-03 | Yes |
| Tap2 | -0.53 | 7.50E-04 | 1.24E-01 | Yes |
| Rif1 | -0.51 | 1.25E-03 | 1.65E-01 | Yes |
| Mycn | 0.50 | 1.65E-03 | 1.82E-01 | No |
| Apoe | 0.48 | 2.25E-03 | 1.93E-01 | No |
| Krt18 | 0.48 | 2.34E-03 | 1.93E-01 | No |

**DEGs of myelo-erythroid restricted vs. multilineage HSC-clone-derived MPP**

| Gene Symbol | log(fold-change) | P value | adj. P value | Up-regulated in multilineage HSC-clone-derived MPP? |
|---|---|---|---|---|
| Flt3 | -1.40 | 1.90E-06 | 1.37E-03 | Yes |
| Dntt | -1.11 | 2.02E-04 | 3.99E-02 | Yes |
| Cebpb | -1.07 | 3.33E-04 | 3.99E-02 | Yes |
| Arl5a | 1.06 | 3.87E-04 | 3.99E-02 | No |
| Eefsec | 0.98 | 1.04E-03 | 1.51E-01 | No |
